# Multilayer control of KaiR1D-autoreceptor function by the auxiliary protein Neto

**DOI:** 10.1101/2025.10.06.680643

**Authors:** Wen-Chieh Hsieh, Tae Hee Han, Rosario Vicidomini, Peter Nguyen, Zheng Li, Mihaela Serpe

**Author notes:** Correspondent Author: Mihaela Serpe Eunice Kennedy Shriver National Institute of Child Health and Human Development National Institutes of Health 35 Convent Drive, Bldg. 35, Room 1C-1016 Bethesda, MD, USA, 20892. These authors contributed equally to this work (Wen-Chieh Hsieh, Tae Hee Han, Rosario Vicidomini).

## Abstract

Kainate-type glutamate receptors and their dedicated auxiliary protein Neto function at both pre- and postsynaptic sites to regulate the activity of synaptic networks. However, attributing specific synaptic functions to Neto and/or kainate receptors is challenging. Here we focus on *Drosophila* KaiR1D receptors, which modulate synaptic transmission at neuromuscular junction, and elucidate the role of Neto in the regulation of autoreceptor activities and neurotransmitter release. We show that Neto-α limits the presynaptic accumulation and function of KaiR1D autoreceptors *in vivo*. Using outside-out patch recordings, we demonstrate that Neto-α modulates the KaiR1D gating properties, slowing desensitization and attenuating the block by intracellular polyamines and extracellular toxins. Neto-α also promotes the axonal distribution of KaiR1D, but this function is not critical for the KaiR1D-dependent regulation of synaptic transmission. Instead, Neto-α increases the charge transfer and likely the KaiR1D-mediated influx of Ca^2+^ in the presynaptic compartment leading to increased neurotransmitter release. Our data demonstrate that Neto-α provides multiple layers of modulation to KaiR1D autoreceptors to ensure proper neurotransmitter release. These findings broaden our view on how auxiliary subunits shape channel gating and subcellular distribution, suggesting that coordinated regulation of receptor function and localization represents an ancestral strategy to safeguard synaptic stability.

## Introduction

Glutamate is the major neurotransmitter at vertebrate excitatory central synapses and at the neuromuscular junction (NMJ) of insects and crustaceans (Hansen *et al*, 2021; Jan & Jan, 1976a; Takeuchi & Takeuchi, 1964). Its actions are mediated primarily via ionotropic glutamate receptors (iGluRs) which were pharmacologically classified in three major classes named α-amino-3-hydroxy-5-methyl-4-isoxazolepropionic acid (AMPA), N-methyl-D-aspartic acid (NMDA), and kainate (KA) receptors (Hansen *et al*., 2021). The cloning of insect iGluRs revealed sequences similar to vertebrate AMPA, KA and NMDA receptors and suggested conserved functional properties between insect iGluRs and their vertebrate counterparts. However, heterologous expression of several *Drosophila* AMPA and KA receptors revealed strikingly different kinetics and agonist binding properties (Han *et al*, 2015; Li *et al*, 2016). Like vertebrate iGluRs, *Drosophila* receptors are modulated by auxiliary proteins. Auxiliary subunits are transmembrane proteins which associate with iGluRs throughout their lifetime and modulate when, where and how the channels function (Jackson & Nicoll, 2011). Genetics and functional reconstitution studies suggest that the iGluRs and their auxiliary subunits evolved together (Orvis *et al*, 2022; Ramos-Vicente &

Bayes, 2020; Ramos-Vicente *et al*, 2021; Stroebel & Paoletti, 2021), so that all AMPA receptors require TARPs (transmembrane AMPAR regulatory proteins) (Jackson & Nicoll, 2011; Li *et al*., 2016), and all KA receptors require Neto (<u>Ne</u>uropillin and <u>To</u>lloid-like) proteins (Copits & Swanson, 2012; Han *et al*., 2015; Tomita & Castillo, 2012). Unlike AMPARs and their suite of auxiliary subunits, it has been challenging to assign specific synaptic functions to Neto in complexes with KARs.

In the vertebrate CNS, KARs function pre- and post-synaptically and serve as both modulators and mediators of synaptic transmission (Contractor *et al*, 2011; Lerma & Marques, 2013). These receptors have been implicated in human diseases ranging from epilepsy to neuropathic pain and migraine (Contractor *et al*., 2011). As such, KAR complexes are attractive pharmacological targets for therapeutic intervention. However, the low abundance of KARs and Neto proteins, the complex array of effects they elicit on neural circuit activity, and the limited availability of pharmacological and histological tools have hindered our progress and understanding. By contrast, insects such as the fruit fly rely on KARs and Neto to mediate and modulate synaptic transmission at the NMJs, an essential synapse for animal viability; in this system, perturbations of KARs or Neto activities produce a wide range of phenotypes, from embryonic paralysis and death to subtle synaptic plasticity deficits (Han *et al*, 2020; Kim *et al*, 2012). Insects expanded their kainate receptor clade and their functional repertoire partly to cover the needs of their glutamatergic NMJs (Li *et al*., 2016). For example, the *Drosophila* genome contains 10 genes coding for kainate receptor subunits, of which at least six are utilized at the NMJ. They form 1) two distinct postsynaptic complexes, GluRIIC, GluRIID and GluRIIE, and either GluRIIA (type-A receptors) or GluRIIB (type-B receptors), that co-exist within individual PSDs and enable NMJ functionality and plasticity (DiAntonio, 2006; Featherstone *et al*, 2005; Marrus *et al*, 2004; Qin *et al*, 2005), and 2) a KaiRID-containing presynaptic complex required for normal neurotransmitter release (Kiragasi *et al*, 2017). We have previously demonstrated that Neto is an obligatory subunit for postsynaptic iGluR NMJ complexes, and that it is required for the recruitment and stabilization of these receptors at synaptic sites (Han *et al*., 2015; Han *et al*, 2024; Kim *et al*., 2012; Kim *et al*, 2015; Ramos *et al*, 2015). In the absence of Neto, iGluRs remain scattered on the muscle surface away from synaptic sites and (*neto^null^*) animals die as paralyzed embryos, unable to hatch into the larval stages of development (Kim *et al*., 2012). We also found that presynaptic Neto facilitates neurotransmitter release (Han *et al*., 2020), a function that requires the KaiR1D autoreceptors (Kiragasi *et al*., 2017), but whether and how Neto shapes the function of KaiR1D-containing complexes remains unknown.

It has been long recognized that KARs, operating as autoreceptors, provide a critical feedback mechanism that modulates neurotransmitter release and ensures stable neuronal network activities (Chittajallu *et al*, 1996; Contractor *et al*., 2011; Lerma & Marques, 2013; MacDermott *et al*, 1999; Schmitz *et al*, 2001). Perturbations to this feedback have been associated with a wide range of neurological, psychiatric and neurodegenerative disorders (Pinheiro & Mulle, 2008). Vertebrate KAR autoreceptor activities have been described at the synapses between mossy fibers (MF) and CA3 pyramidal cells in the hippocampus (MF-CA3 synapses), where they contribute to several forms of synaptic plasticity (Pinheiro *et al*, 2007). Furthermore, presynaptic KARs and Neto1 guide synaptic connectivity in the developing hippocampus (Orav *et al*, 2017). At the *Drosophila* NMJ, KaiR1D autoreceptor and Neto have been both positively implicated in the control of glutamate release (Han *et al*., 2020; Kiragasi *et al*., 2017). The *Drosophila neto* gene codes for two isoforms (Neto-α and Neto-β) that share 1) highly conserved extracellular domains, including two extracellular CUB (for C1r/C1s–Uegf–BMP) domains and an LDLa motif, and 2) one transmembrane pass, but have completely different intracellular domains of 206 and 351 residues, generated by alternative splicing (Ramos *et al*., 2015). The two isoforms have different *in vivo* distributions and functional roles, with Neto-β, the predominant postsynaptic isoform, providing a dynamic scaffold that recruits both iGluRs and PSD components and sculpts the PSD composition, and Neto-α functioning in both pre- and postsynaptic compartments (Han *et al*., 2020; Kim *et al*., 2012; Kim *et al*., 2015; Ramos *et al*., 2015). Postsynaptic Neto-α limits the PSD size, whereas presynaptic Neto-α, like KaiR1D, is required for normal basal transmission and homeostatic plasticity (Han *et al*., 2020).

Here, we investigate the role of Neto-α in modulating the properties of presynaptic KaiR1D receptors and KaiR1D-dependent neurotransmitter release. We found that *Drosophila* Neto-α limits KaiR1D *in vivo* activity and controls the presynaptic distribution and function of KaiR1D receptor channels. Using outside-out patch-clamp recordings and fast ligand application, we examined the biophysical properties of KaiR1D receptors expressed in HEK293T cells and found that Neto-α modulates KaiR1D gating properties, slowing desensitization and attenuating their block by intracellular polyamines and extracellular toxins. When expressed in primary rat hippocampal neurons, *Drosophila* Neto-α promotes KaiR1D axonal entry and presynaptic accumulation; in this system, Neto-α and KaiR1D colocalize in the proximity of active zones marked by the presynaptic scaffold protein Bassoon, suggesting that Neto-α stabilizes KaiR1D at presynaptic locations. Our structure-function studies provide mechanistic insights on functional regulation and demonstrate that Neto-α is critical for increasing the KaiR1D-mediated charge transfer in the synaptic compartment, facilitating the neurotransmitter release.

## Results

### Neto-α limits the function of KaiR1D *in vivo*

To examine the role of Neto-α in the regulation of KaiR1D function, we first tested whether selective overexpression of KaiR1D in motor neurons could restore the basal transmission levels in the absence of Neto-α. To facilitate detection of KaiR1D receptors, we generated a C-terminal Flag tagged variant (details in Materials and Methods). When overexpressed in motor neurons (using a motor neuron specific driver, *BG380-Gal4*), KaiR1D-Flag fully rescued the lethality and basal neurotransmission deficits of *KaiR1D^null^* mutants. We recorded from muscle 6 (abdominal segment 3) in third instar larvae and found that the evoked junction potentials (EJPs) decreased by 40% in the absence of KaiR1D (to 19.94 ± 1.82 mV from 34.15 ± 0.97 mV in control, p<0.0001) (Fig. 1A-B, Supplemental Table 1). Overexpression of KaiR1D-Flag in motor neurons rescued the basal neurotransmission deficits of *KaiR1D^null^* to wild-type levels (33.21 ± 0.76 mV) (Fig. 1C, quantified in Fig.1E-F). This indicates that addition of the Flag tag does not perturb KaiR1D *in vivo* function. Similar to *KaiR1D^null^* mutants, the EJPs are reduced in the absence of Neto-α [Fig. 1F and (Han *et al*., 2020)]; overexpression of KaiR1D-Flag in this genetic background cannot restore the basal neurotransmission, which remains closer to the levels recorded at *neto-*α*^null^* NMJs (18.97 ± 0.24 mV vs 22.01 ± 0.67 in *neto-*α*^null^*, p= 0.6437) (Fig. 1D-F and Supplemental Table 1). This result indicates that Neto-α limits KaiR1D-dependent basal neurotransmission.

**Figure 1.**
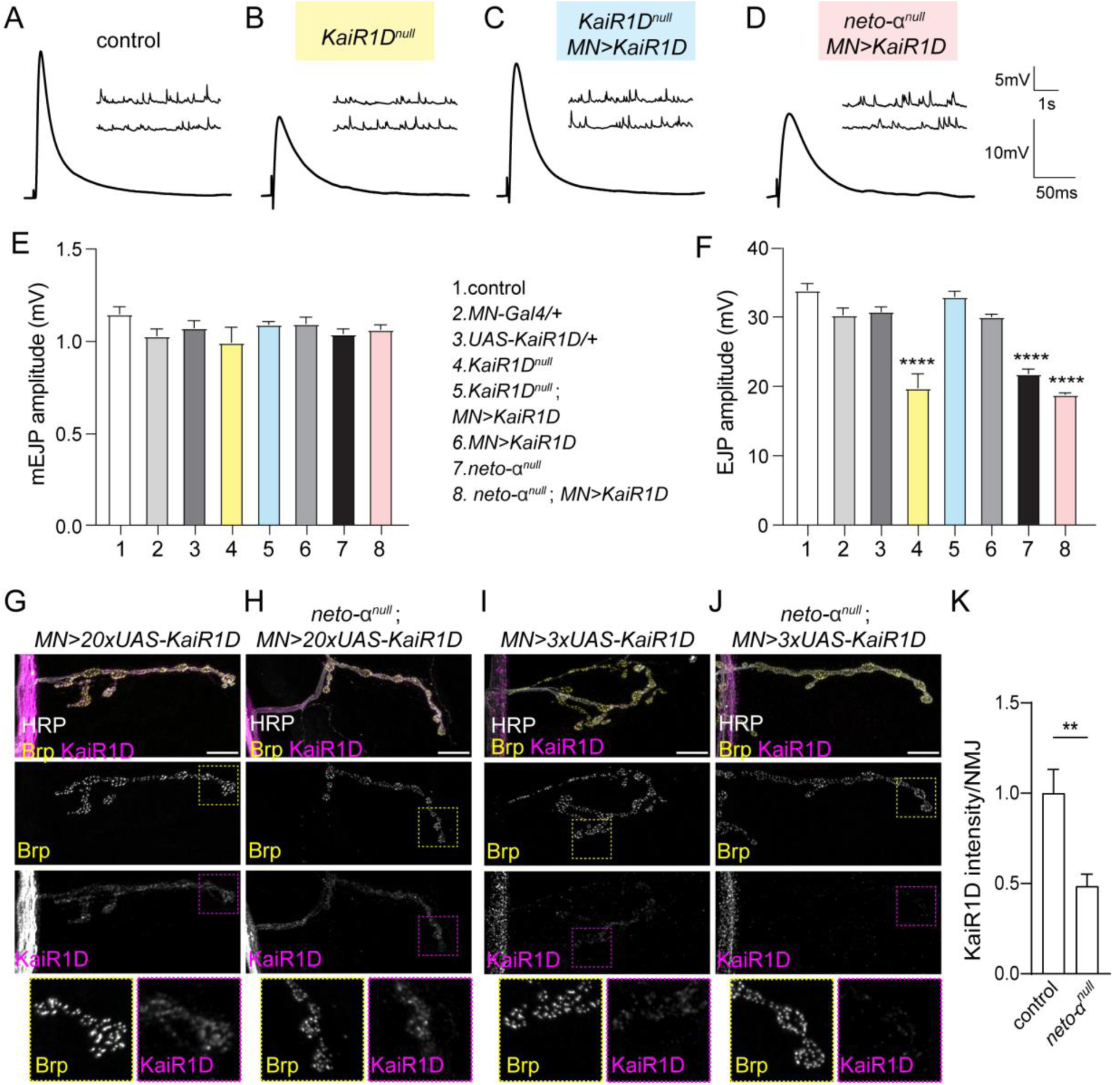
**Neto-α limits the function and distribution of KaiR1D *in vivo*** (A-D) Representative traces for mEJP and EJP recordings from muscle 6 (abdominal segments 3-4) of the indicated genotypes. (E-F) Quantification of mEJP (E) and EJP (F) amplitudes. Overexpression of KaiR1D in motor neurons could not overcome the absence of Neto-α. (G-J) Confocal images of third instar larva NMJ stained for KaiR1D (magenta), Brp (yellow) and HRP (white). High level KaiR1D expression (*20xUAS* transgene) enables delivery/detection of KaiR1D at synaptic terminals with (G) or without Neto-α (H). However, limited levels of KaiR1D expression (*3xUAS* transgene) can only detected at presynaptic terminals in the presence (I) but not in the absence of Neto-α (J), quantified in (K). Statistical analyses for electrophysiology recordings are done with one-way ANOVA with post hoc Tukey’s multiple comparisons; for immunohistochemistry we used unpaired two-tailed Mann-Whitney tests. Data are represented as mean ± SEM.****p < 0.0001; **p < 0.01. Scale bars: 10 μm.

Overexpression of KaiR1D-Flag in an otherwise wild-type background did not induce any changes in basal neurotransmission: The EJPs measured were comparable to wild-type or driver alone controls (Fig. 1F). It is important to note that the large EJP differences recorded here occurred in the absence of any significant changes in the amplitude of miniature excitatory junction potentials (mEJPs), which represent the postsynaptic response to the spontaneous release of single synaptic vesicles (Fig. 1E). This indicates that genetic manipulations of KaiR1D primarily disrupt presynaptic neurotransmitter release leaving the postsynaptic receptor fields largely unaffected.

The neurotransmission deficits observed at *neto-*α*^null^* NMJs were not caused by abnormal expression or trafficking of KaiR1D-Flag in the absence of Neto-α. For these initial experiments we used a transgene with 20 Gal4 binding elements (*20xUAS*), which ensures high levels of expression in both control and *neto-α^null^* motor neurons (Fig. 1G-H, Supplemental Fig. S1). Also, KaiR1D-Flag trafficked normally to the synaptic terminals and accumulated around the presynaptic active zone marked by the presynaptic scaffold Bruchpilot (Brp), the fly homolog of the vertebrate active zone protein ELKS/CAST (Kittel *et al*, 2006; Wagh *et al*, 2006). Since overloading the presynaptic terminals with KaiR1D cannot compensate for loss of Neto-α activities, our results indicate that Neto-α controls the *in vivo* function of KaiR1D-containing autoreceptors.

Neto proteins have been previously implicated in the axonal distribution of mammalian KARs (Orav *et al*., 2017). In flies as in vertebrate systems, presynaptic KARs and Neto-α are low abundance proteins and multiple attempts to generate antibodies that detect endogenous proteins failed. Instead, to examine whether *Drosophila* Neto-α influences the presynaptic localization of KaiR1D, we titrated down the levels of KaiR1D-Flag expression (using a transgene with only three Gal4 binding elements, *3xUAS-KaiR1D-Flag*) (Fig. 1I-K, Supplemental Fig. S1). In this setting, KaiR1D was detectable in motor neuron bodies in both control and *neto-α^null^* mutants but was significantly reduced at *neto-α^null^* NMJs (to 49 ± 7% of the control levels, p= 0.0034). By contrast, the active zone scaffold Brp equally labeled the presynaptic terminals in both genotypes, providing an internal control. The drastic reduction of KaiR1D signals at *neto-α^null^* NMJs indicates that *Drosophila* Neto-α promotes presynaptic localization of KaiR1D.

### *Drosophila* Neto proteins modulate KaiR1D gating kinetics and channel conductance

To investigate the role of Neto in the modulation of KaiR1D function, recombinant KaiR1D was transiently transfected in HEK293T cells with or without Neto variants and incubated for three days at 30°C prior to outside-out patch recordings. Since KaiR1D has low affinity to glutamate (Li *et al*., 2016), we examined the gating properties in response to rapid application of 10 mM glutamate. When transfected alone, KaiR1D assembled as functional homomeric ion channels with relatively low success rate; 28% patches (n= 42 of 151) elicited glutamate-gated currents, suggesting a low level of KaiR1D expression and/or surface delivery [Fig. 2 and (Li *et al*., 2016)]. When KaiR1D was transfected with Neto proteins, the success rate doubled (to 65% for KaiR1D/Neto-α, n= 44 of 68). Rapid application of 10 mM glutamate to outside-out patches from HEK cells transfected with various KaiR1D/Neto combinations revealed differential effects of Neto variants (Supplemental Table 2). For macroscopic currents recorded from complexes of KaiR1D alone, the 10%-90% rise time was 237.1 ± 10.1 µs, with a deactivation time constant (τ_off_) 0.39 ± 0.03 ms (Fig. 2A). KaiR1D/Neto complexes had similar rise times and deactivation time constants (Supplemental Table 2). Longer applications of glutamate (100 ms) revealed a role for Neto proteins in the desensitization of KaiR1D channels, with the decay best fit by the sum of two exponential functions: KaiR1D: τ_fast_ 1.07 ± 0.06 ms, τ_slow_ 12.72 ± 1.61 ms, A_fast_ 89.61 ± 2.61%; KaiR1D/Neto-α, τ_fast_ 4.30 ± 0.35 ms, τ_slow_ 17.21 ± 1.45 ms, A_fast_ 75.89 ± 3.98%; and KaiR1D/Neto-β, τ_fast_ 3.90 ± 0.58 ms, τ_slow_ 17.58 ± 2.21 ms, A_fast_ 91.12 ± 3.44 % (Fig 2H). To better describe the receptor decay time, a weighted tau (τ_w_) was calculated (see Methods). As previously reported, KaiR1D has very fast desensitization, with the weighted decay time τ_w_ 2.31 ± 0.39 ms; Neto-α slows down the desensitization of KaiR1D channels 3-fold (τ_w_ 7.09 ± 0.34 ms, P< 0.0001). Neto-β has a smaller, 2-fold impact (τ_w_: 5.15 ± 0.73 ms, p= 0.0009). Since Neto-β cannot traffic to presynaptic locations (Han *et al*., 2020) and remains confined to the somato-dendritic compartment, its effect on KaiR1D desensitization could be relevant for KaiR1D postsynaptic complexes, for example in the adult eye (Karuppudurai *et al*, 2014). Notably, the desensitization time constants for KaiR1D/Neto-ΔCTD complexes (ΔCTD, deletion of C-terminal domain), τ_fast_ 2.94 ± 0.44 ms, and τ_w_ 4.33 ± 0.47 ms, were comparable to values measured KaiR1D/Neto-β complexes (Fig. 2E, G-H), indicating that a “minimal Neto”, which retains the highly conserved extracellular and transmembrane domains, but lacks any intracellular parts, also has ∼2-fold impact on the desensitization of KaiR1D (τ_w_ 4.33 ± 0.47 ms vs τ_w_ 2.31 ± 0.39 ms for KaiR1D alone).

**Figure 2.**
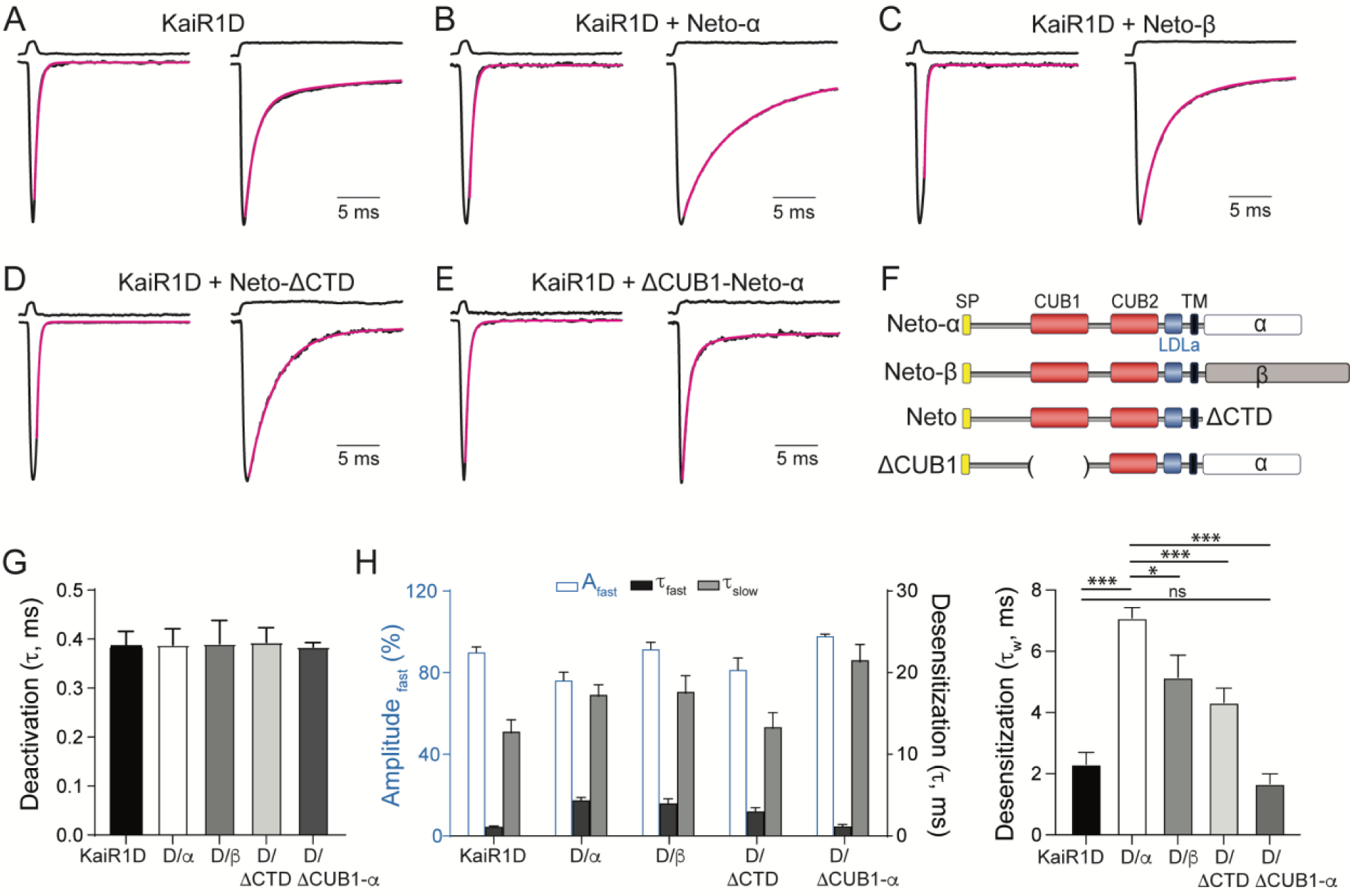
**The effect of Neto variants on KaiR1D deactivation and desensitization** (A - E) Responses to 10 mM glutamate applied for 1 ms (left traces) and 100 ms (right traces) to outside-out patches from cells transfected with KaiR1D alone (A) or together with Neto variants, Neto-α (B), Neto-β (C), Neto-ΔCTD (D) or ΔCUB1-Neto-α (E). Black lines show the average of 15-20 responses from one patch normalized and aligned to the peak. Red lines show decay of the responses fitted with the with the sum of one (deactivation) or two (desensitization) exponential functions; open tip junction currents measured at the end of the experiments are shown at the top of each panel. The holding potential was - 60mV for all recordings. (F) Diagram of the Neto variants utilized. (G-H) Summary graph for deactivation (G) and desensitization time constants (H) for various KaiR1D/Neto complexes; the A_fast_ (%) component for desensitization is shown in blue. Statistical analyses are done with one-way ANOVA with post hoc Tukey’s multiple comparisons. Data are represented as mean ± SEM. ***p < 0.001; *p < 0.05; ns, p > 0.05.

Interestingly, deletion of the CUB1 domain of Neto-α induced even faster desensitization of KaiR1D/ΔCUB1-Neto-α channels (τ_fast_: 1.12 ± 0.22 ms, τ_slow_: 21.46 ± 2.19 ms; A_fast_ 97.57± 0.93%, τ_w_: 1.68 ± 0.32 ms, n= 8, Fig. 2F-H). This result is consistent with previous reports that CUB1 domain of vertebrate Neto proteins stabilizes the structure and function of recombinant kainate receptor complexes in a heterologous expression system (Li *et al*, 2019; Zhang *et al*, 2006). Also, recent Cryo-EM studies revealed that vertebrate Neto2 engages GluK2 via two anchor points, one between Neto2-CUB1 and GluK2-ATD and another in the transmembrane region (He *et al*, 2021). The deletion of CUB1 removes one of these anchor points and is predicted to disrupt the Neto-α/KaiR1D interaction; this may affect the CUB2-LBD interface, influencing the gating properties of KaiR1D-containing complexes.

To further determine the effect of Neto-α on KaiR1D channel properties, we performed non-stationary fluctuation analysis (Fig. 3A–B, quantified in 3C–F). For this analysis we used a custom Verify_R pipeline (Supplementary file) and examined 50-98 trials/patch from six outside-out patches for each receptor complex. Estimates of single-channel conductance showed a two-fold increase in the conductance of KaiR1D/Neto-α channels (79.6 ± 11.3 pS) compared with KaiR1D alone (38.3 ± 6.1 pS, p= 0.0094) (Fig. 3. A-D). By contrast, Neto-α induces a small, but significant increase in the open probability, 0.95 ± 0.1 for KaiR1D/Neto-α versus 0.89 ± 0.1 for KaiR1D alone (p= 0.0143) (Fig. 3E). The conductance of KaiR1D channels is in the range of those measured for mammalian KARs, *i.e.* 27 pS for GluK2 homomeric channels (Zhang *et al*, 2009); however, different from KaiR1D/Neto-α, co-expression of NETO2 did not affect the conductance of GluK2 channels but had a marked impact (4-fold increase) on their open probability. These results demonstrate that Neto-α modulates the biophysical properties of KaiR1D receptors and underscore species-specific differences between the effects of Neto proteins on kainate receptor unitary properties in flies versus mammals.

**Figure 3.**
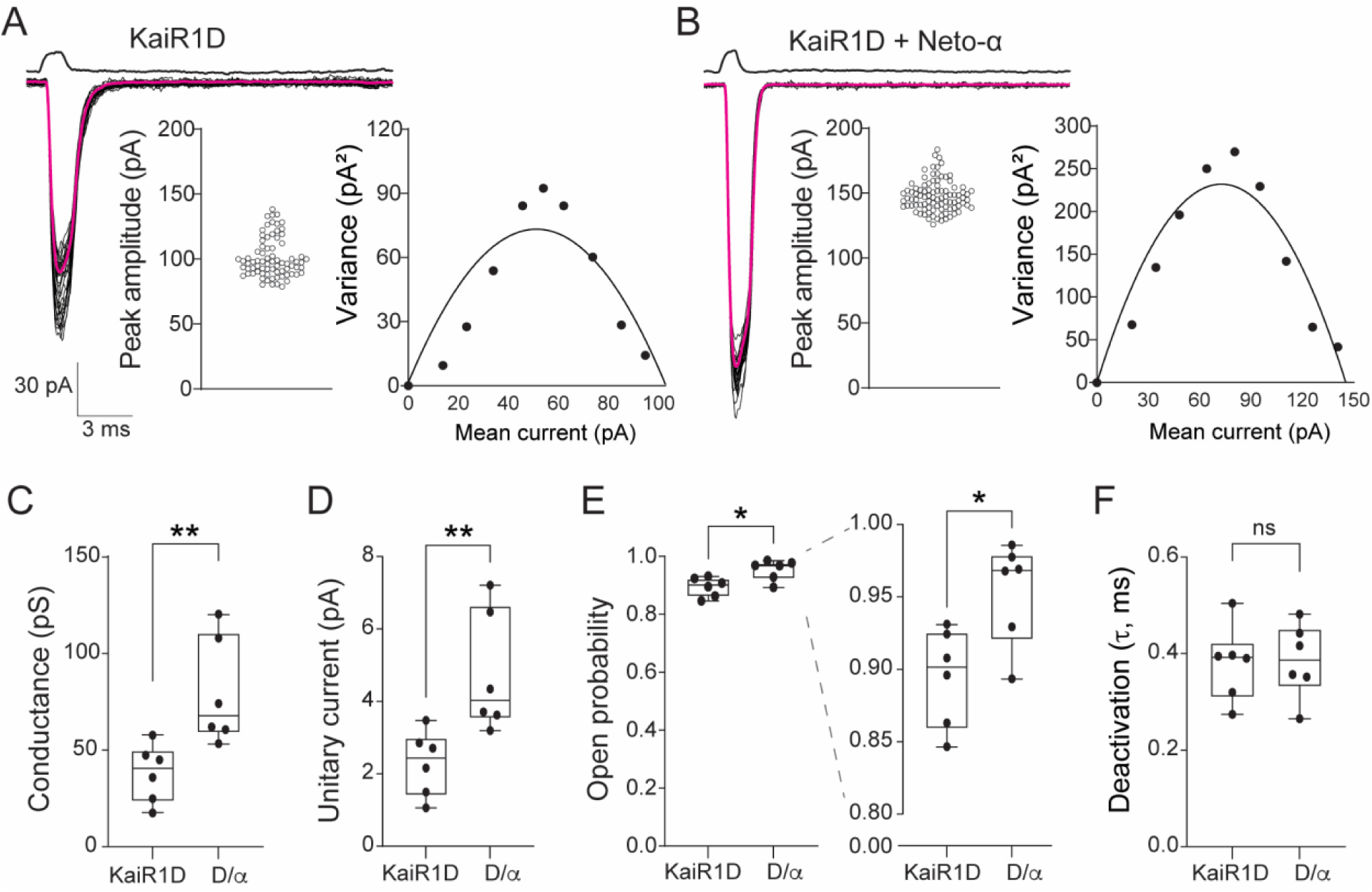
Non-stationary analysis of variance of responses to 10mM glutamate applied for 1 ms to outside-out patches from HEK cells transfected with KaiR1D receptor complexes. (A - B) Superimposed individual responses are shown in black; the average current in magenta; open junction current at the end of the experiments are shown at the top. The holding potential was −60 mV for all recordings. The insets in each panel show the peak amplitude for all trials (left) and the current-variance relationship (right) fit with the function… (C-F) Conductance (C), open probability (D), unitary currents (E) and deactivation time constants (F) determined by variance analyses of responses for six patches for both KaiR1D and KaiR1D/ Neto-α channel complexes as indicated, showing responses for individual patches. Statistical analyses are done with one-way ANOVA with post hoc Tukey’s multiple comparisons. Data are represented as mean ± SEM. **p < 0.01; *p < 0.05; ns, p>0.05.

### Neto variants modulate block of desensitization by Concanavalin A

The lectin Concanavalin A (Con A) attenuates the desensitization of KARs for a wide variety of species, including native vertebrate KARs (Huettner, 1990; Wong & Mayer, 1993), *Drosophila* recombinant receptors (Han *et al*., 2015) and AvGluR1 from the primitive eukaryote *Adineta vaga* (Lomash *et al*, 2013). In prior work, we found that pretreatment with 0.6 mg/ml Con A for 10 min attenuates the desensitization to glutamate for KaiR1D expressed in *Xenopus* oocytes (Li *et al*., 2016). In the absence of Con A, application of 10 mM glutamate for 100 ms to outside-out patches of HEK cells containing KaiR1D receptors produced robust desensitization (94.8 ± 1.27%); pretreatment with of Con A reduced desensitization (73.4 ± 5.03%) with a 5-fold increase in amplitude of the glutamate-evoked steady-state for the entire duration of glutamate application (Fig. 4A). KaiR1D/Neto-α receptor channels showed almost complete (92.4 ± 2.05%) albeit slower desensitization, while pretreatment with Con A practically abolished desensitization (Fig. 4B). This Con A-induced block of desensitization does not require the intracellular domain of Neto-α: KaiR1D/Neto-ΔCTD channels showed a similar robust block of desensitization (95.5 ± 0.56%) with strong steady-state currents for the entire duration of glutamate application (Fig. 4C, quantified in D).

**Figure 4.**
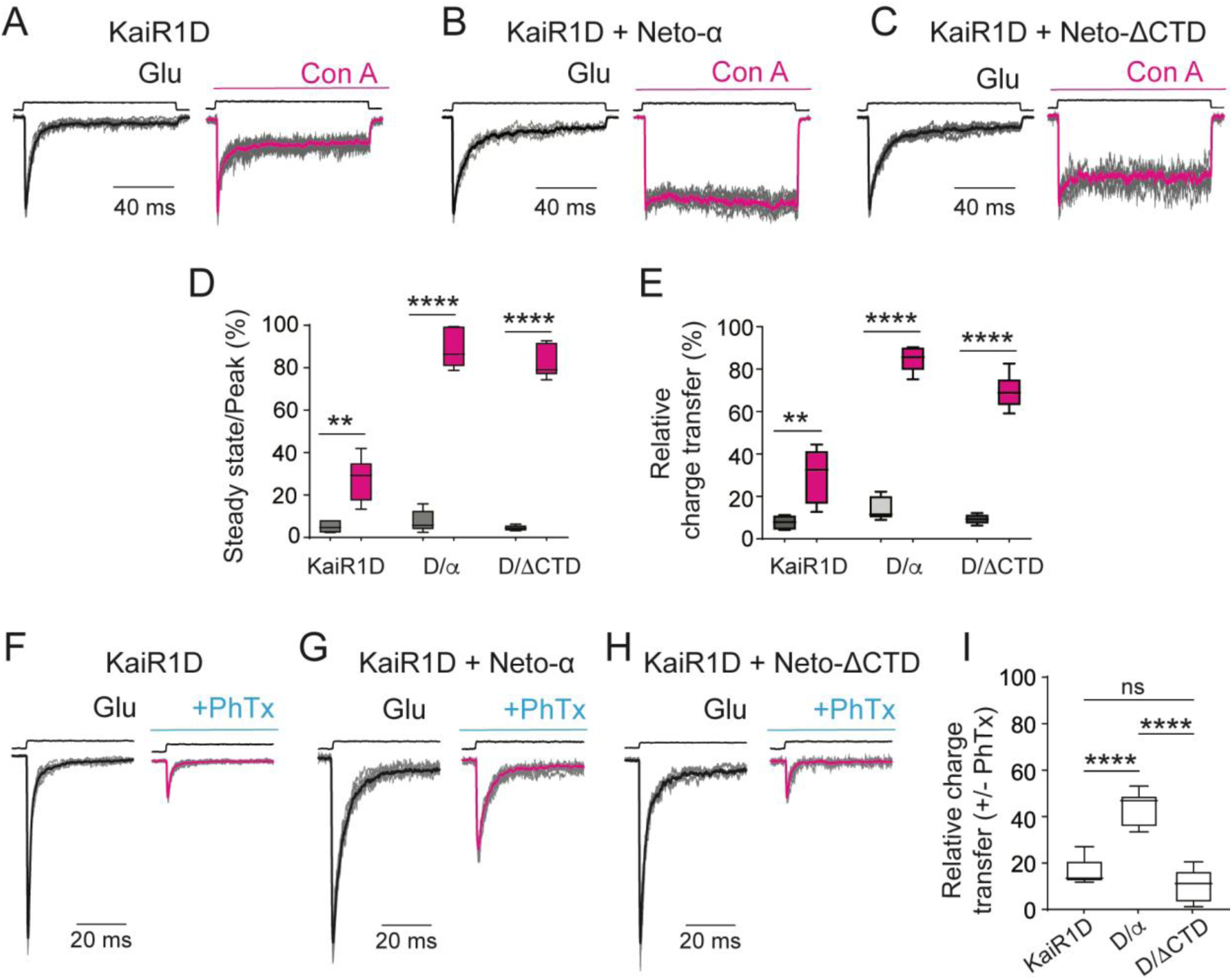
The effect of Concanavalin A and philanthotoxin on KaiR1D receptor complexes. (A-C) Responses (gray) evoked at a holding potential of −60mV by rapid application of 10 mM glutamate for 100 ms to outside-out patches with superimposed average from cells expressing KaiR1D alone (A) or together with Neto-α (B) or Neto-ΔCTD (C), before (left panels) and after (right panels) treatment with 0.6 mg/ml Con A for 10 min. Black and red lines show the average of 35-60 responses from one patch; open tip junction currents measured at the end of the experiments are shown at the top of each panel. The holding potential was −60mV for all recordings. Con A induces profound desensitization of KaiR1D, which is completely blocked by both Neto-α and Neto-ΔCTD. (D) Average ratio of steady-state current/peak current from outside-out patches expressing KaiR1D-containing complexes before (black) and after (red) 10 min treatment with 0.6 mg/ml Con A. (E) Charge transfer measured by the integration of 18-27 trials for each channel divided by their maximum value (peak amplitude multiplied by the time of glutamate application, 100 ms). (F-H). Responses (gray) of glutamate-evoked currents induced by rapid application of 10 mM glutamate for 100 m to outside-out patches from HEK cells expressing KaiR1D-containing complexes and recorded at a holding potential of −60mV. Superimposed average from before (black) and after (red) application of 1 μM philanthotoxin-433 (PhTx) are shown. (I) Bar graph showing the relative charge transfer (%) of KaiR1D-containing channels induced by 1 μM PhTx. Statistical analyses are done with one-way ANOVA with post hoc Tukey’s multiple comparisons. Data are represented as means ± SEM. ****p < 0.0001; ns, p > 0.05.

Calculation of the charge transfer by integration of raw data in response to application of glutamate for 100 ms revealed substantial differences between KaiR1D alone and KaiR1D in complexes with either Neto-α or Neto-ΔCTD. Relative to the total current during the 100 ms application of glutamate, the charge transfer was 7.75 ± 1.39% (n= 5), 14.4 ± 2.18% (n= 6) and 9.28 ± 0.95% (n= 5) for KaiR1D, KaiR1D/Neto-α and KaiR1D/Neto-ΔCTD, with a large increase for patches treated with Con A, relative charge transfer 30.10 ± 5.10% (n= 6), 84.60 ± 2.46% (n= 6) and 69.36 ± 3.23% (n= 6) (Fig. 4E).

### Neto-α attenuates the external philanthotoxin block of KaiR1D

Ca^2+^-permeable KARs are blocked by extracellular philanthotoxin (PhTx) (Bahring *et al*, 1997), a polyamine toxin derived from wasp venom (Eldefrawi *et al*, 1988; Karst & Piek, 1991). Indeed, external PhTx blocked the response of KaiR1D to glutamate in recordings from *Xenopus* oocytes, with several minutes required for full recovery from the block (Li *et al*., 2016). We tested whether Neto influences the PhTx block by applying 1 µM PhTx to outside-out patches, while recording responses of receptor channels to 100 ms pulses of 10 mM glutamate. PhTx caused a substantial block of all KaiR1D receptor channel complexes analyzed (Fig. 4F-H, quantified in I). Before application of PhTx, the mean charge transfer in response to 100 ms applications of 10 mM was: KaiR1D, 2.65 ± 1.81 pC; KaiR1D/Neto-α, 8.89 ± 3.46 pC; KaiR1D/Neto-ΔCTD, 6.15 ± 3.13 pC (Figure 4H). Inhibition by PhTx developed slowly, τ_onset_ 88 s for KaiR1D alone, 154 s for KaiR1D/Neto-α and 90 s for KaiR1D/Neto-ΔCTD (Supplemental Fig. S2). At equilibrium, PhTx markedly reduced the charge transfer by 83.7 ± 2.78% (n= 5) and 89.6 ± 3.26% (n= 5) of control, for KaiR1D and KaiR1D/Neto-ΔCTD channels respectively. For KaiR1D/Neto-α channels, the block at equilibrium was weaker, with the charge transfer reduced by 55.84 ± 3.02% (n= 6) compared to control.

The slow onset reduction in charge transfer by PhTX suggests a progressive decrease in the number of channels opening in response to glutamate in the presence of PhTx (Supplemental Fig. S2A-C). In addition, the rate of decay of the response to glutamate, which in control conditions results from the onset of desensitization, increases most likely due to a combination of desensitization and open channel block by PhTx. Indeed in the presence of PhTx, the kinetics of desensitization/channel block was substantially faster, as estimated from the average response to 15-30 applications of glutamate recorded at steady state, τ_w_ KaiR1D 1.80 ± 0.56 ms before and 0.91 ± 0.30 ms after PhTx (n = 5); τ_w_ KaiR1D/Neto-α 6.96 ± 0.65 ms vs 3.82 ± 0.42 ms (n = 6); τ_w_ KaiR1D/Neto-ΔCTD 4.02 ± 0.29 ms vs 2.09 ± 0.36 (n = 5). Most of this change was due to an increase in the fast component of decay of the response to glutamate (Supplemental Table 3). The slow onset of PhTx block (Supplemental Fig. S2) is consistent with the requirement of channel opening for toxin entry and likely reflects the low open probability and rapid desensitization of KaiR1D.

Together, our recordings demonstrate that *Drosophila* Neto-α modulates the gating kinetics of KaiR1D channels, increases the mean channel conductance and alters the pharmacological profile.

### Neto-α colocalizes with KaiR1D and promotes KaiR1D axonal distribution

In *Drosophila* larvae lacking Neto-α, we observed reduced KaiR1D localization at motor neuron terminals (Fig. 1I-K). However, additional unknown modulators might contribute to KaiR1D entering the axonal compartment and trafficking to presynaptic terminals in larval motor neurons. To eliminate the contribution of such unknown modulators and test for a direct role for Neto in KaiR1D axonal distribution, we set up an *ex vivo* experiment, transfecting recombinant *Drosophila* KaiR1D and Neto-α in 15-day-in-vitro (DIV15) cultured primary rat hippocampal neurons and examining their subcellular distribution four days later (DIV19); Venus was also introduced to visualize the transfected neurons. Primary hippocampal neurons lack detectable Neto-1 and −2 mRNAs and have been previously utilized to study the subcellular distribution of mammalian KARs (Copits *et al*, 2011; Palacios-Filardo *et al*, 2016). This *ex vivo* system should eliminate the contribution of other *Drosophila* KaiR1D modulators and permit the identification of Neto-α molecular determinants required for KaiR1D presynaptic distribution.

We found that, when expressed by itself, KaiR1D-Flag was marginally detectable in the axon albeit it accumulated as clear puncta in the somato-dendritic compartment (Fig. 5A-C). In contrast, Neto-α positive puncta of various sizes were visible in both dendritic and axonal compartments (Fig. 5D-F). The HA tag did not influence Neto-α subcellular localization as tagged and untagged Neto-α variants showed similar distributions when expressed in primary hippocampal neurons (Supplemental Fig. S3). Co-expression of KaiR1D and Neto-α together did not change the distribution of Neto-α to neurites but had a significant effect on the presence of KaiR1D in the axonal compartment (Fig. 5G-I). The axon initiation segment, labeled with Ankyrin G, localized within 30 μm from the cell soma; consequently, we examined the axonal compartment starting from >100 μm away from the soma. We found that increased numbers of KaiR1D positive puncta decorate the axon and colocalize with Neto-α signals. This suggests that Neto-α promotes KaiR1D axonal localization. To confirm this apparent increase of axonal KaiR1D distribution and to account for variable transfection efficiency among different neurons/experiments, we quantified and normalized the density of KaiR1D puncta in axons versus dendrites in the absence or presence of Neto-α (Fig. 5J). Neto-α increased KaiR1D puncta density observed along the axons, from 0.38 ± 0.06, n= 30 cells, for KaiR1D alone to 1.02± 0.14, n= 32, (p<0.0001) for KaiR1D/Neto-α. These results are consistent with our *in vivo* observations that Neto-α promotes KaiR1D presynaptic distribution (Fig.1I-K).

**Figure 5.**
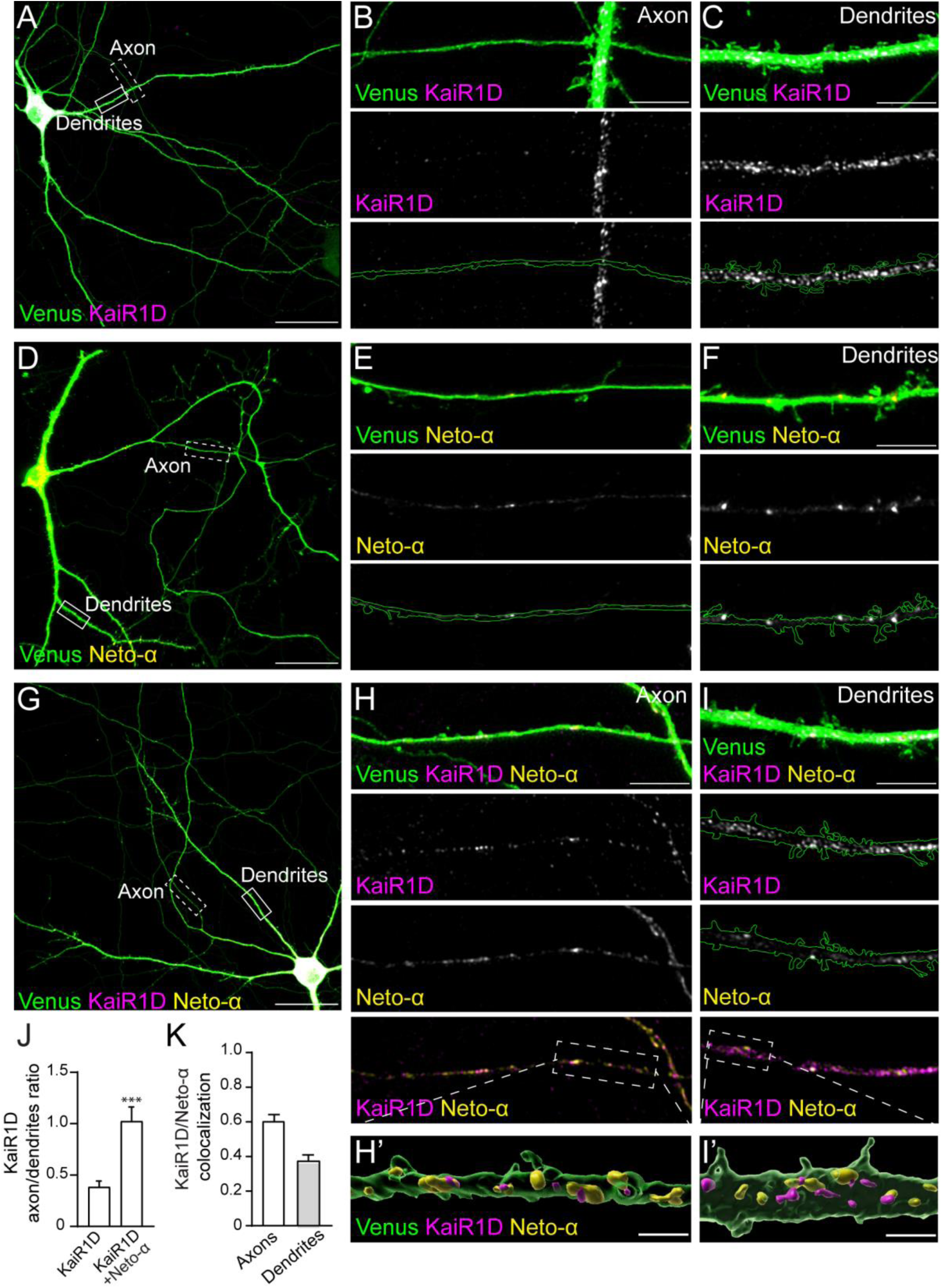
**Neto-α promotes KaiR1D axonal distribution.** (A-I) Confocal images of primary rat hippocampal neurons transfected (DIV15) and stained (DIV19) for KaiR1D (magenta), Venus (green) and Neto-α (yellow). Low magnification views (A, D and G) capture the cell body as well as part of the axon and dendrites, shown at higher magnification in details (B-C, E-F and H-I). (J) Quantification of KaiR1D puncta density in axon vs. secondary dendrites. (K) Colocalization between KaiR1D and Neto-α in axons and secondary dendrites. (L-M) 3D reconstitution of subcellular localization of KaiR1D and Neto-α along the axon (L) or in the dendrites (M). Data are represented as mean ± SEM. Indicated P values are from unpaired two-tailed Mann-Whitney tests. ***p < 0.001. Scale bars: 40 µm (A, D and G), 5 µm (B, C, E, F, H and I) and 1 µm (H’ and I’).

There was limited colocalization between KaiR1D and Neto-α in the dendrites (Fig. 5K and I). This may reflect the nature of these proteins, since both *Drosophila* KaiR1D and Neto-α likely lack the protein features/interaction motifs required for recruitment and stabilization at vertebrate excitatory synapses. For example, unlike Neto-α, both mammalian Neto proteins have PDZ binding motifs (Tomita & Castillo, 2012). Alternatively, at 37°C, the fly proteins may display inappropriate interactions or post-translational modifications that disrupt their own association and/or synaptic recruitment in this *ex vivo* mammalian system. We tested the distribution of KaiR1D and Neto-α in hippocampal primary neurons maintained at 30°C and found no change compared to 37°C. Since Neto family proteins have highly divergent intracellular domains that have been implicated in the synaptic recruitment of glutamate receptors (Copits & Swanson, 2012; Lomash *et al*, 2017; Straub & Tomita, 2012; Tang *et al*, 2011), we favor the explanation that *Drosophila* KaiR1D ± Neto-α complexes lack the ability to interact with mammalian synaptic scaffolds.

By contrast, a large proportion of axonal KaiR1D puncta colocalized with Neto-α signals (584 out of 889, n= 32 cells) (Fig. 5K-H’). These results suggest that KaiR1D and Neto-α form complexes that traffic together and/or interact at specific locations along the axons. In flies as well as in vertebrate systems, Neto proteins and KARs stabilize each other at synaptic sites (Kim *et al*., 2012; Zhang *et al*., 2009). Thus, it is possible that *Drosophila* KaiR1D and Neto-α colocalize at presynaptic sites when co-expressed in primary hippocampal neurons. In mammals, presynaptic locations are marked by Bassoon, a scaffolding protein involved in the active zone organization (tom Dieck *et al*, 1998). Indeed, without Neto-α, we rarely observed accumulation of KaiR1D-positive puncta in the vicinity of Bassoon immunoreactivities (45 out of 703 puncta analyzed). When co-transfected with Neto-α, the number of KaiR1D positive puncta colocalized with Bassoon slightly increased (87 out of 1170) but more importantly, puncta positive for both KaiR1D and Neto-α were significantly enlarged (Supplemental Fig. S4). These observations are consistent with stabilization of KaiR1D/Neto-α complexes at presynaptic sites. Together, our results indicate that Neto-α promotes KaiR1D axonal distribution and colocalizes with KaiR1D in the axonal compartment; however, KaiR1D and Neto-α may travel largely independent of each other to the dendrites.

### The Neto-α cytoplasmic domain promotes KaiR1D axonal delivery

Structural analyses of vertebrate KARs demonstrated that Neto proteins bind via two anchor points, one between the NTD and CUB1 domains and one within the transmembrane region (He *et al*., 2021). If KaiR1D binds to and travels together with Neto-α, then removing one of the anchor points may impact KaiR1D axonal distribution. We tested this possibility by expressing KaiR1D together with ΔCUB1-Neto-α in primary hippocampal neurons. Like Neto-α, ΔCUB1-Neto-α distributed as puncta throughout the neurons (Fig. 6A-C). Surprisingly, KaiR1D axonal delivery was not diminished in the presence of ΔCUB1-Neto-α and instead appeared enhanced in comparison to KaiR1D co-expressed with Neto-α (Fig. 6D-F, quantified in G). This indicates that the CUB1 domain is dispensable for KaiR1D axonal delivery. Removal of CUB1 domain also did not impact KaiR1D and ΔCUB1-Neto-α colocalization, which resembled the KaiR1D/Neto-α distribution pattern, namely prominent colocalization along the axons and sparse in the dendritic compartment (Fig. 6H). In contrast, a C-terminal truncated Neto, Neto-ΔCTD, which lacks any intracellular domain, distributed to both axons and dendrites when transfected alone, but could no longer promote KaiR1D axonal delivery in co-expression experiments (Fig. 6I-N). Neto-ΔCTD did not impact KaiR1D distribution along the dendritic shaft. In addition, Neto-ΔCTD partially colocalized with KaiR1D puncta within the dendrites and with the rare KaiR1D signals observed along the axons (Fig. 6H). Together these results indicate that the intracellular domain of Neto-α is critical for the distribution of KaiR1D to axons.

**Figure 6.**
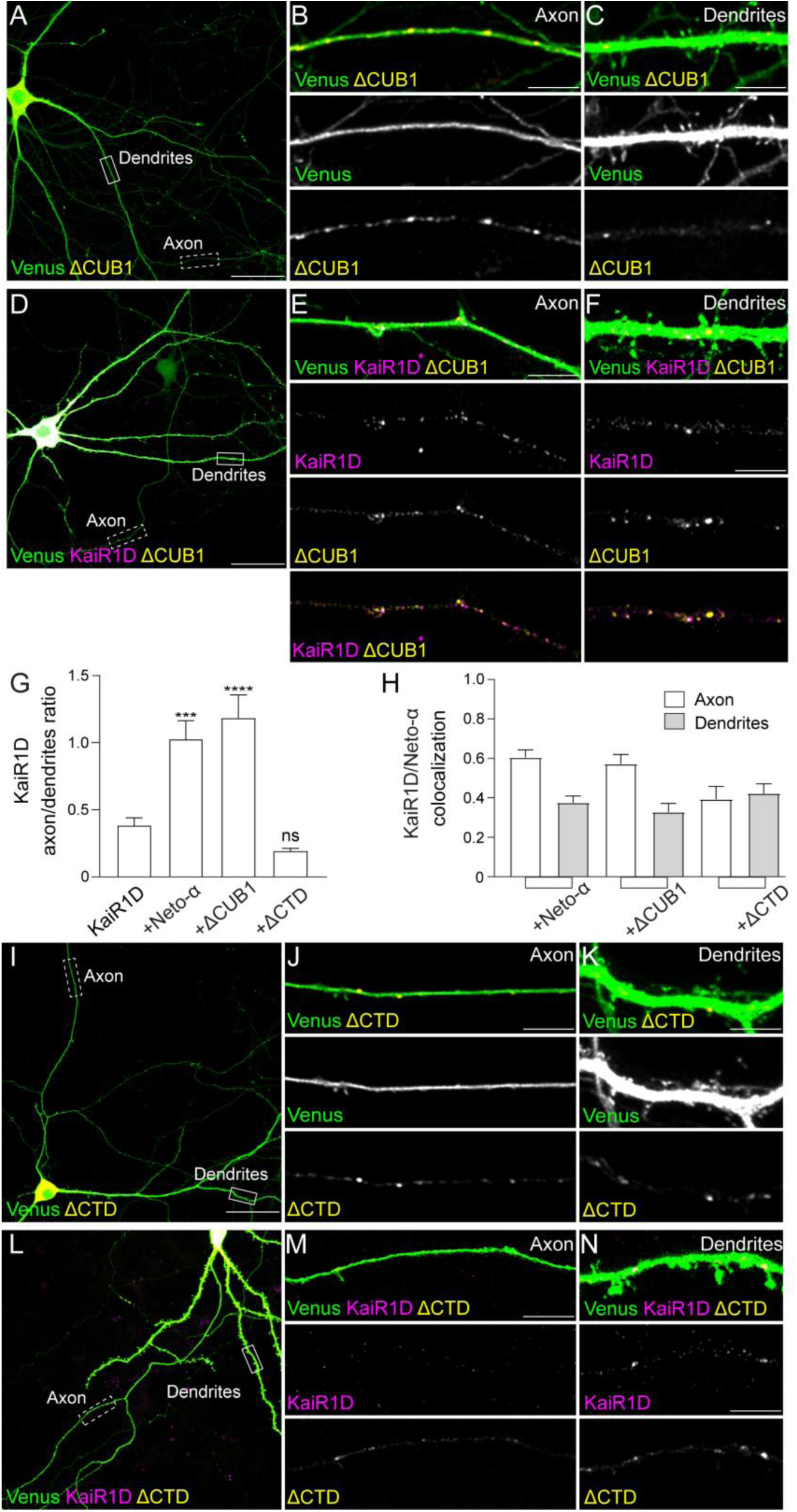
**Neto-α cytoplasmic domain enables KaiR1D axonal delivery** (A-F) Confocal images of primary rat hippocampal neurons transfected (DIV15) and stained (DIV19) for KaiR1D (magenta), Venus (green) and ΔCUB-Neto-α (yellow). Global views (right panels) capture the cell body as well as part of the axon and dendrites, shown at higher magnification in details (middle and left panels). (G) Quantification of KaiR1D puncta density in axon vs. secondary dendrites of hippocampal neurons transfected with KaiRID alone (0.38 ± 0.06, n= 30) or together with Neto-α (1.02 ± 0.14, n= 32, p= 0.0083), Neto-α-ΔCUB1 (1.19 ± 0.17, n= 27, p= 0.0009) and Neto-ΔCTD (0.58 ± 0.17, n= 30, p= 0.7606) as indicated. (H) Colocalization between KaiR1D and Neto-α variants in axons and secondary dendrites. (I-N) Confocal images of primary rat hippocampal neurons transfected and labeled for KaiR1D (magenta), Venus (green) and Neto-α-ΔCUB1 (yellow). Neto-ΔCTD cannot promote KaiR1D axonal delivery. Data are represented as mean ± SEM. Indicated P values are from one-way ANOVA with Karuskal-Wallis multiple comparison test. ****p < 0.0001; ***p < 0.001; ns, p > 0.05. Scale bars: 40 µm (A, D and G) and 5 µm (axons and dendrites details).

### Neto-α CUB1 domain is essential for the NMJ function *in vivo*

Our *in vitro* and *ex vivo* studies indicate that Neto-α modulates both the function and axonal distribution of KaiR1D-containing autoreceptors. Are any of these Neto-α functions essential for normal KaiR1D-dependent neurotransmitter release *in vivo*? To address this, we examined whether the Neto-α CUB1 and CTD domains are required for larval NMJ function using *in vivo* rescue experiments. To facilitate a direct comparison among Neto variants, we used phiC31 integrase-mediated transformation to introduce all *neto* transgenes at the same chromosomal location (the attP docking site vk14) (Venken *et al*, 2006), and ensure similar levels of expression (see Materials and Methods).

As previously reported, EJPs recorded at larval muscle 6 (abdominal segments 3 and 4) were reduced by half in absence of Neto-α (from 34.18 ± 0.56 mV in control to 22.01 ± 0.67 mV in *neto-α^null^*, p<0.0001) (Fig. 7A, Supplemental Table 4). Overexpression of full-length Neto-α in motor neurons rescued the EJPs at *neto-α^null^* NMJs (to 34.08 ± 1.37 mV Fig. 7B). In contrast, overexpression of ΔCUB1-Neto-α did not rescue basal neurotransmission at *neto-α^null^* NMJs; the EJPs recorded in these animals were 24.37 ± 1.57 mV, closer to EJPs levels measured in *neto-α^null^* (22.01 ± 0.67 mV, p= 0.7978) and clearly different from control (34.15 ± 0.97 mV, p< 0.0001) (Fig. 7C). This difference was not due to different protein levels or impaired traffic to presynaptic terminals since all Neto variants concentrated robustly at synaptic boutons (Supplemental Fig. S5). Instead, the inability of ΔCUB1-Neto-α to rescue basal neurotransmission in *neto-α^null^* mutants indicates that this Neto variant cannot sustain KaiR1D function *in vivo*. This may reflect the extremely fast desensitization kinetics of KaiR1D/ΔCUB1-Neto-α channels (Fig. 2) and/or the inability of ΔCUB1-Neto-α to properly engage KaiR1D receptor complexes in the absence of the CUB1-mediated anchor point (He *et al*., 2021). If ΔCUB1-Neto-α engages KaiR1D inefficiently, then overexpression of ΔCUB1-Neto-α in motor neurons should have minimal impact on neurotransmission; on the other hand, if ΔCUB1-Neto-α binds to KaiR1D and does not slow desensitization of these channels *in vivo*, then excess ΔCUB1-Neto-α should compete with endogenous Neto-α for KaiR1D and reduce the EJPs. We found that neuronal overexpression of ΔCUB1-Neto-α in an otherwise wild-type background reduces the basal transmission levels (to 26.33 ± 1.18 mV, p< 0.0001) (Fig. 7F, Supplemental Table 4). This result suggests that excess ΔCUB1-Neto-α displaces endogenous Neto-α from presynaptic KaiR1D/Neto-α complexes; the accumulation of fast desensitizing KaiR1D/ΔCUB1-Neto-α complexes then reduces neurotransmitter release.

**Figure 7.**
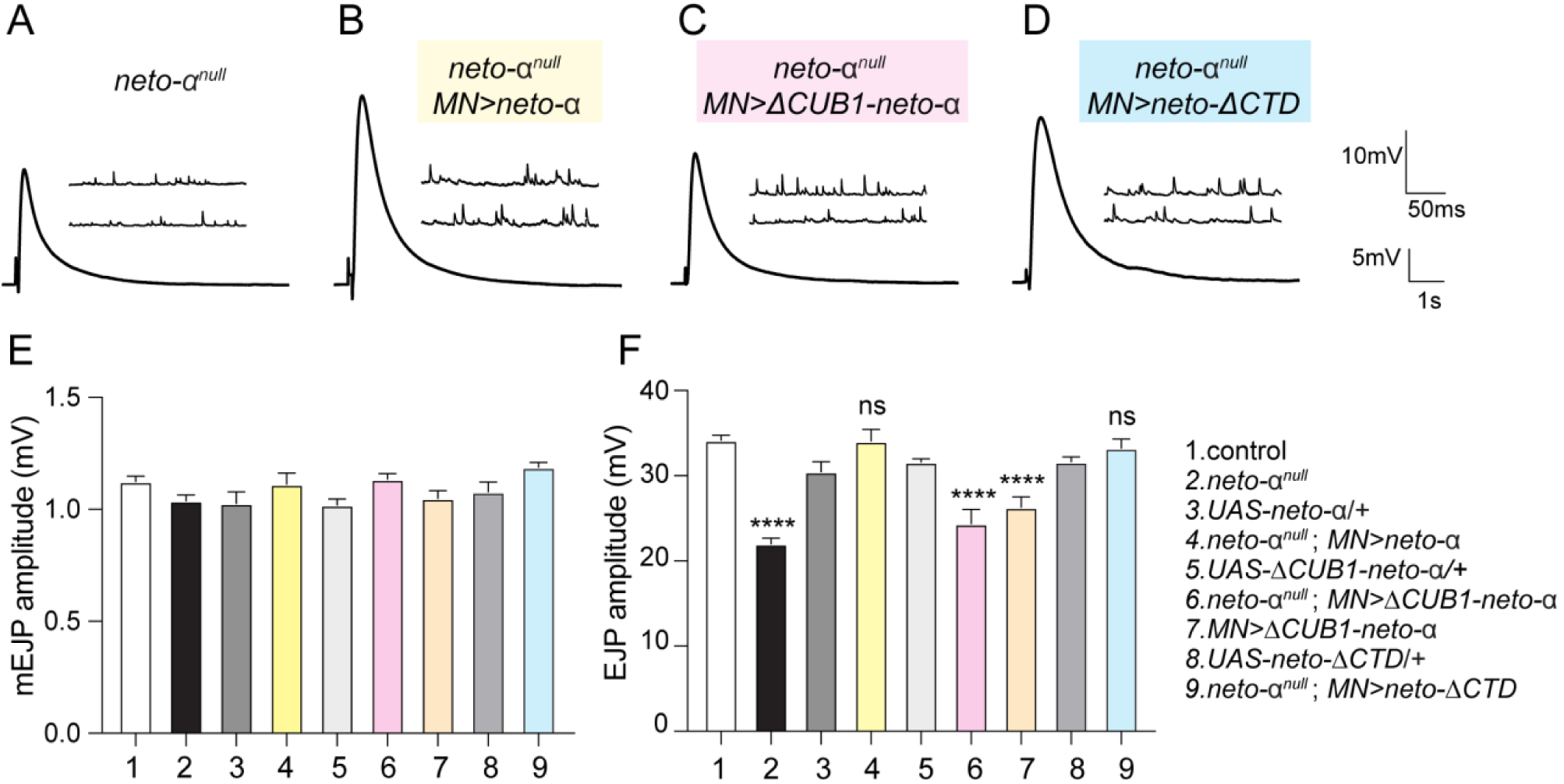
**Modulation of KaiR1D function is key for normal basal neurotransmission *in vivo*** (A-D) Representative traces for mEJP and EJP recordings from muscle 6 (abdominal segments 3-4) of the indicated genotypes. (E-F) Quantification of mEJP and EJP amplitudes. Overexpression of Neto-α variants that slow down KaiR1D desensitization rescues basal neurotransmission levels at *neto-*α*^null^* NMJs, whereas Neto-α-ΔCUB1 does not, even though it promotes KaiR1D axonal delivery. Statistical analyses are done with one-way ANOVA with post hoc Tukey’s multiple comparisons. Data are represented as mean ± SEM.****p < 0.0001; ns, p > 0.05

In contrast, neuronal overexpression of Neto-ΔCTD fully rescued the EJPs at *neto-α^null^* NMJs (to 33.24 ± 1.08 mV, Fig. 7D). This finding is consistent with our previous observation that overexpression of either full-length Neto-α-GFP or Neto-ΔCTD-GFP rescues neurotransmission deficits at *neto-α^null^* NMJs (Han *et al*., 2020). These results suggest that the role of Neto-α in promoting KaiR1D axonal distribution is either not essential or redundant for NMJ function *in vivo*. Other interacting proteins may contribute to KaiR1D axonal delivery and/or stabilization at presynaptic terminals *in vivo*. By contrast, excess ΔCUB1-Neto-α which sustains KaiR1D axonal delivery but accelerates channel desensitization, is unable to rescue basal neurotransmission at *neto-α^null^* NMJs. Together these data demonstrate that Neto-α-dependent KaiR1D desensitization is a key element for maintaining normal basal neurotransmission at this synapse.

## Discussion

Here we combined powerful *Drosophila* genetics with outside-out patch recordings and *ex vivo* analyses to examine the role of the auxiliary protein Neto in the function and axonal distribution of KaiR1D-containing presynaptic receptors. We found that Neto-α is required for KaiR1D-dependent basal neurotransmission through multiple mechanisms. Neto-α promotes axonal distribution of KaiR1D receptors and appears to stabilize them at synaptic sites. Neto-α also modulates the gating properties of KaiR1D receptor channels, slowing desensitization threefold and doubling the channel conductance. Different Neto-α domains control distinct activities: the CUB1 domain is critical for Neto-mediated modulation of channel properties but not for axonal distribution, whereas the intracellular domain of Neto-α influences both KaiR1D gating properties and axonal delivery. Our study captures a Neto-enabled multidimensional control of KaiR1D receptor channels distribution and function that safeguards the stability and reliability of neurotransmission.

### Neto-α shapes KaiR1D-dependent basal neurotransmission

The roles of Neto-type proteins in the biology of presynaptic KARs has been difficult to parse out. The low levels of both KARs and Neto proteins as well as the heterogeneity and subsynaptic distribution of KARs both confounded the issue. Presynaptic KARs can act as autoreceptors or heteroreceptors and function to both facilitate and depress neurotransmitter release in various regions of the brain [reviewed in (Contractor *et al*., 2011; Pinheiro & Mulle, 2008)]. Several presynaptic kainate receptor/Neto complexes with different subunit compositions have been identified at mammalian synapses (Orav *et al*., 2017; Wyeth *et al*, 2017). By contrast, only one presynaptic KaiR1D-containing autoreceptor and one Neto-type protein, Neto-α, function to facilitate neurotransmitter release at the *Drosophila* NMJ (Han *et al*., 2020; Kiragasi *et al*., 2017). If Neto-α functions as a true auxiliary subunit for KaiR1D receptors, then its presence should be required for normal KaiR1D-mediated activities in the larval motor neurons. Indeed, we found that KaiR1D, even when in excess, cannot sustain normal neurotransmitter release in the absence of Neto-α (Fig. 1D). In previous studies, we found that KaiR1D probably forms Ca^2+^-permeable ion channels with low affinity for glutamate (EC50 10.7 mM) (Li *et al*., 2016). The low Glu affinity is similar to the levels observed for postsynaptic KARs at larval NMJ (Han *et al*., 2015) and likely reflects the channels adaptation to the high Glu concentration in the hemolymph, the blood fly counterpart, which bathes all internal tissues (Piyankarage *et al*, 2008). A KaiR1D transgene with a Gln 604 to Arg mutation, a position in the pore loop equivalent to the Q/R site which undergoes RNA editing in vertebrate iGluRs (Higuchi *et al*, 1993; Kohler *et al*, 1993), does not restore normal basal transmission at *KaiR1D^null^* NMJs (Kiragasi *et al*., 2017). This suggested that KaiR1D facilitates glutamate release by increasing presynaptic Ca^2+^ influx. Additional KaiR1D-mediated contributions, such as metabotropic signaling (Lerma & Marques, 2013) could be possible but are yet to be unexplored.

We found that Neto-α controls KaiR1D autoreceptor function by 1) promoting the axonal distribution of KaiR1D at presynaptic sites, and 2) by modulating KaiR1D channel properties and gating kinetics. The physiological significance of Neto-α-dependent KaiR1D modulation, including reduced desensitization, increased channel conductance, and increased open probability, would enhance KaiR1D-mediated charge transfer, including Ca^2+^ influx, in the presynaptic motor neuron. Apart from its role in receptor trafficking, Neto-α’s ability to potentiate KaiR1D currents is essential for normal basal transmission at fly NMJ: When overexpressed, KaiR1D can reach motor neuron terminals but does not facilitate synaptic transmission in the absence of Neto-α (Fig. 1D-E), presumably because rapid desensitization limits the duration of Ca^2+^ influx. By contrast, overexpression of mouse GluK1c rescues the neuronal development deficits of *Neto1^-/-^* mice, even though NETO1 promotes axonal delivery of GluK1c (Orav *et al*., 2017). It is however possible that such developmental deficits, which include impaired synaptogenesis, reduced excitability and synchronization of neuronal populations, require smaller GluK1c-mediated currents or could be compensated by other mechanisms.

The prominent roles for *Drosophila* Neto proteins in iGluR biology likely follow the expansion of the kainate receptor clade in arthropods (Li *et al*., 2016; Orvis *et al*., 2022): *Drosophila* genome codes for 10 kainate-type receptors, six of which are essential for the formation of NMJ and hence for fly viability (Featherstone *et al*., 2005; Marrus *et al*., 2004; Qin *et al*., 2005). We identified *Drosophila neto* as an essential gene required for the synaptic recruitment and function of postsynaptic NMJ iGluRs (Kim *et al*., 2012; Ramos *et al*., 2015); in fact, the functional reconstitution of postsynaptic NMJ iGluRs was only possible in the presence of either Neto isoform (Han *et al*., 2015; Han *et al*., 2024). Conversely, the AMPAR and NMDAR clades have been somewhat contracted during the evolution of insects (Moroz *et al*, 2021). The *Drosophila* genome has only two AMPARs and two NMDARs that appear to be encoded by non-essential genes, as several homozygous viable genetic null alleles have been generated for each of these loci. By contrast, mammals rely on AMPARs and NMDARs for neuronal development and function while their five KARs are not essential for organism viability (Xu *et al*, 2017). AMPA, kainate and NMDA, the canonical ligands used to classify mammalian receptors, elicit very different responses in insects or other non-vertebrate systems (Li *et al*., 2016). For example, we found that postsynaptic NMJ KARs expressed in *Xenopus* oocytes respond to glutamate, but not AMPA or kainate (Han *et al*., 2015); also, the fly AMPA-type receptor GluR1A responds to kainate, but not AMPA (Li *et al*., 2016). Glutamate, kainate and quisqualate activate KaiR1D, whereas NMDA and D-AP5, both agonists for vertebrate NMDARs, inhibit KaiR1D (Li *et al*., 2016). Despite these major differences in ligand recognition among glutamate receptor clades and their biological relevance, the KAR modulatory functions of Neto auxiliary proteins have been remarkably conserved across species.

### Structure-function studies provide mechanistic insights on functional regulation

Our structure-function studies provide mechanistic insights on the Neto-dependent functional regulation of KaiR1D. Both fly Neto-α and Neto-β isoforms slow KaiR1D desensitization, by 3- and 2.2-fold respectively, but do not impact receptor deactivation (Fig. 2). Notably, the difference between Neto-β and Neto-ΔCTD is not significant, highlighting a role for Neto-α intracellular domain. It has been postulated that Neto modulation occurs through interactions with the KARs LBD, perhaps by influencing the dimer interface which is key to receptor desensitization (Chaudhry *et al*, 2009). Additional contributions could come from motifs that couple ligand binding to receptor gating (Griffith & Swanson, 2015; Horning & Mayer, 2004; Poulsen *et al*, 2019). We found that the CUB1 motif is absolutely required for Neto modulation of KaiR1D desensitization (Fig. 2). Similarly, CUB1-deleted NETO1 and NETO2 mammalian proteins have no apparent effect on GluK2 desensitization (Li *et al*., 2019). The CUB1 domain serves as one of the anchor points for the GluK2/Neto-2 interactions in both inhibited and desensitized states (He *et al*., 2021). Thus, ΔCUB1-Neto-α likely associates with KaiR1D TM1, the second anchor point of KAR/Neto complexes, but strays away from the KaiR1D ligand binding domain; this conformation renders ΔCUB1-Neto-α unable to augment KaiR1D function. Instead, we found that excess ΔCUB1-Neto-α functions as a dominant negative *in vivo*, in larval motor neurons, presumably because it displaces the endogenous Neto-α from KaiR1D/Neto-α presynaptic complexes and speeds the desensitization of KaiR1D channels below the threshold required to facilitate neurotransmitter release (Fig. 7F).

The effect of Neto-α on KaiR1D kinetics could be partly explained by the behavior of KaiR1D-mediated currents at single channel level, which show that Neto-α enhances the channel conductance two-fold (Fig. 3C) and induces a modest increase in open probability (Fig. 3E). This is different from mammalian Neto proteins, which increase the channel open probability by 2-4-fold and have no effect on channel conductance (Tomita & Castillo, 2012). To our knowledge, *Drosophila* KaiR1D is the only example where the channel conductance of a kainate receptor is modulated by a Neto protein. This behavior resembles that of AMPARs, for which association with most auxiliary proteins, including TARPs and Cornichons, increases the mean channel conductance (Jackson & Nicoll, 2011).

We have previously reported that recombinant KaiR1D channels, but not KaiR1D(Q604R), expressed in *Xenopus* oocytes are blocked by PhTx (Li *et al*., 2016). In the present study, we found that Neto-α, through its intracellular domain, attenuates the PhTx-induced block and slows down its onset (Figs. 4 and S2). It was previously demonstrated that the interaction of cytoplasmic polyamines with KARs is influenced not only by Q/R editing but also by a conserved negative charge at the +4 site, near the entrance to the channel (Panchenko *et al*, 1999). NETO1 and NETO2 decrease the inward rectification of GluK2 receptors through positively charged residues located right after their transmembrane domains (Fisher & Mott, 2012); the favored model is that the access of positively charged polyamines to the pore of the channel is altered by these positively charged residues of NETO1 and NETO2. This model may also explain the attenuation of the PhTx-induced block for *Drosophila* KaiR1D/Neto-α complexes since KaiR1D retains the highly conserved +4 negative charge and Neto-α-CTD carries several groups of positive residues. In the future, it will be important to map the specific Neto-α motifs responsible for this attenuation to clarify the underlying mechanism.

### Neto-α facilitates KaiR1D axonal distributions

KaiR1D has a prolonged M4 transmembrane helix and no additional C-terminal tail, lacking any obvious C-terminal regulatory or protein interactions domains. Instead, KaiR1D trafficking and subcellular distribution may be regulated via other domains and intracellular loops as well as by its interaction with Neto-α. The ATD domains of vertebrate KARs have been implicated in the surface expression and synaptic recruitment of these receptors (Duan *et al*, 2018; Matsuda, 2017); also, phosphorylation of Neto-2 appears to restrict GluK1 synaptic targeting (Lomash *et al*., 2017). *Drosophila* has two Neto isoforms with different intracellular domains (Ramos *et al*., 2015), but only Neto-α or Neto-ΔCTD variants can enter the axonal compartment, while Neto-β remains confined to the somato-dendritic region [Fig. S5 and (Han *et al*., 2020)]. It was proposed that somato-dendritic vesicles are sorted and prevented from entry into the axon at the pre-axonal exclusion zone, located in the axon hillock (Farias *et al*, 2015); by contrast, coupling to an axonal kinesin overcomes such vesicle exclusion and allows for their axonal entry and trafficking to presynaptic locations. Therefore, the intracellular domain of Neto-β may contain a motif that prevents binding to axonal carriers, directing this isoform exclusively to somato-dendritic locations. The different subcellular distribution of Neto-α and Neto-β indicates that Neto-α modulates all presynaptic KAR complexes in the fly nervous system.

When expressed in rat hippocampal neurons, KaiR1D traffics to the cell membrane and localizes to the dendrites but rarely in the axonal compartment (Fig. 5). If this limited KaiR1D axonal delivery reflects the effect of endogenous Neto proteins, then knock down of rat NETO1 and NETO2 should eliminate KaiR1D presence in axons; however, this experiment is not technically possible within the relatively long time required for the expression of recombinant KaiR1D in primary hippocampal neurons. Also, it is highly unlikely that endogenous Netos contribute to KaiR1D axonal delivery in rat hippocampal neurons, as NETO1 and NETO2 mRNAs could not be detected in hippocampal cells in culture (Palacios-Filardo *et al*., 2016). Co-expression of KaiR1D with Neto-α increases the frequency and the size of KaiR1D-positive puncta within the axons, suggesting that Neto-α stabilizes and/or promotes KaiR1D clustering at presynaptic locations. These findings highlight the value of using mammalian primary neurons to investigate the subcellular distribution of *Drosophila* KARs. A similar Neto-dependent axonal distribution was reported for mouse GluK1c expressed in primary hippocampal neurons (Orav *et al*., 2017); like KaiR1D, GluK1c partially colocalized with synaptophysin-positive vesicle release sites. This remarkable resemblance between the distribution of mouse GluK1c and *Drosophila* KaiR1D recombinant receptors expressed in primary neurons underscores the high conservation among KARs and Neto proteins.

Based on our results, we propose that KaiR1D carries an intracellular motif that restricts axonal entry and favors its distribution to somato-dendritic vesicles. Neto-α binds to and masks this motif via its CTD, facilitating the coupling of KaiR1D/Neto-α-containing vesicles to axonal kinesin(s) and increasing the axonal distribution of KaiR1D/Neto-α complexes. The intracellular domain of Neto-α is rich in putative phosphorylation sites, raising the possibility that Neto-α provides dynamic regulation for KaiR1D axonal delivery and localization.

Our study uncovers multiple roles for Neto-α in shaping the excitatory transmission: Neto-α secures the deployment of KaiR1D receptor complexes to axonal release sites and strengthens the signaling of individual channels (conductance, gating). Such multilayered control, choreographed through the functions of a single auxiliary subunit, ensures that KaiR1D activities are both potent and precisely localized. This Neto-integrated control mechanism parallels the multifunctionality known for AMPAR auxiliary proteins and indicates that Neto imparts an equally sophisticated regulatory complexity to KARs. In the future, it will be important to examine a role for Neto as a signaling hub that receives information about the network status and adjusts KAR synaptic levels and channel properties, fine-tuning the amplitude of the synaptic output. In evolutionary terms, the presence of this multilayered regulation of a KAR autoreceptor in *Drosophila* implies that coupling receptor function and localization may represent an ancient and conserved strategy to guarantee reliable neurotransmission.

## Materials and Methods

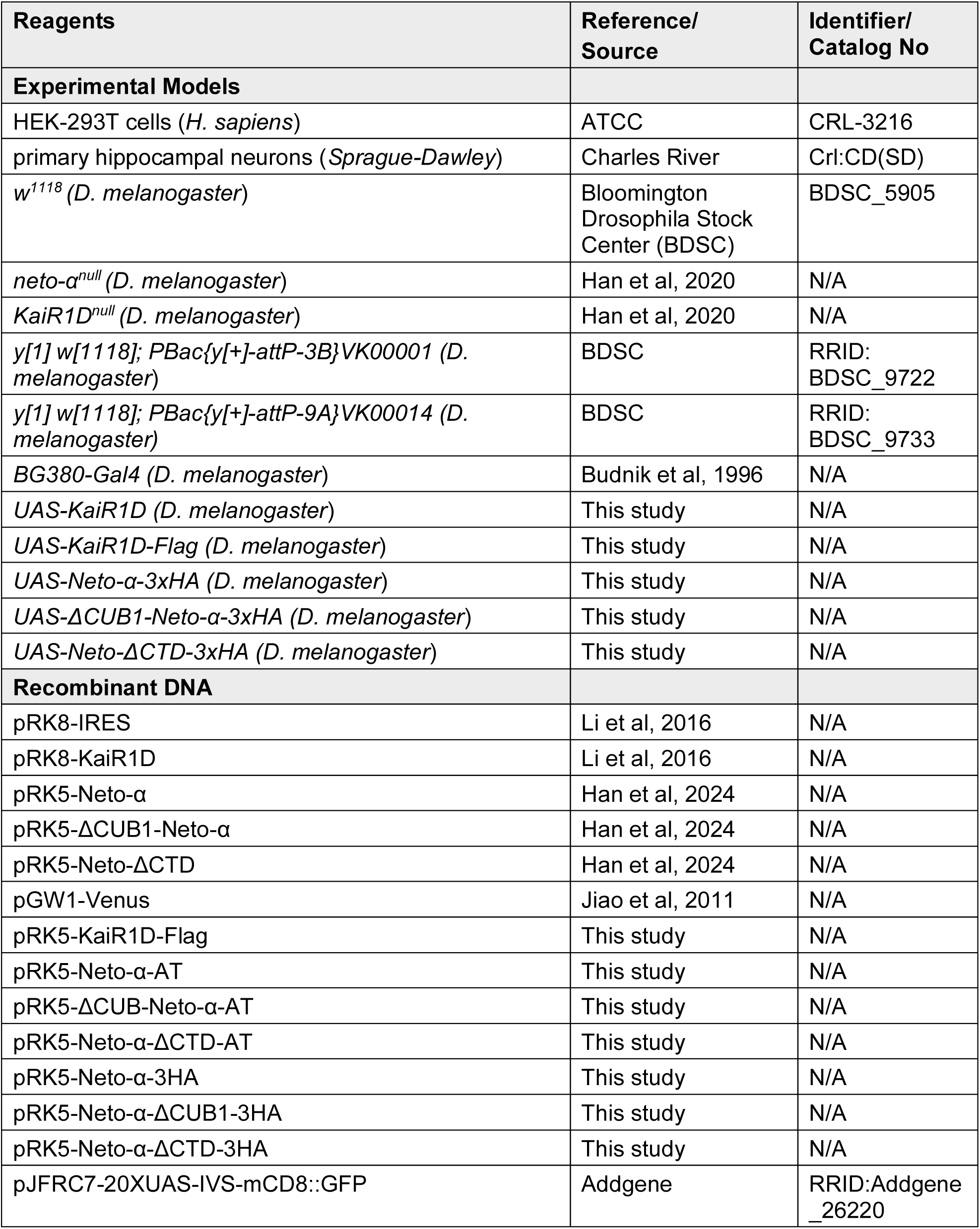
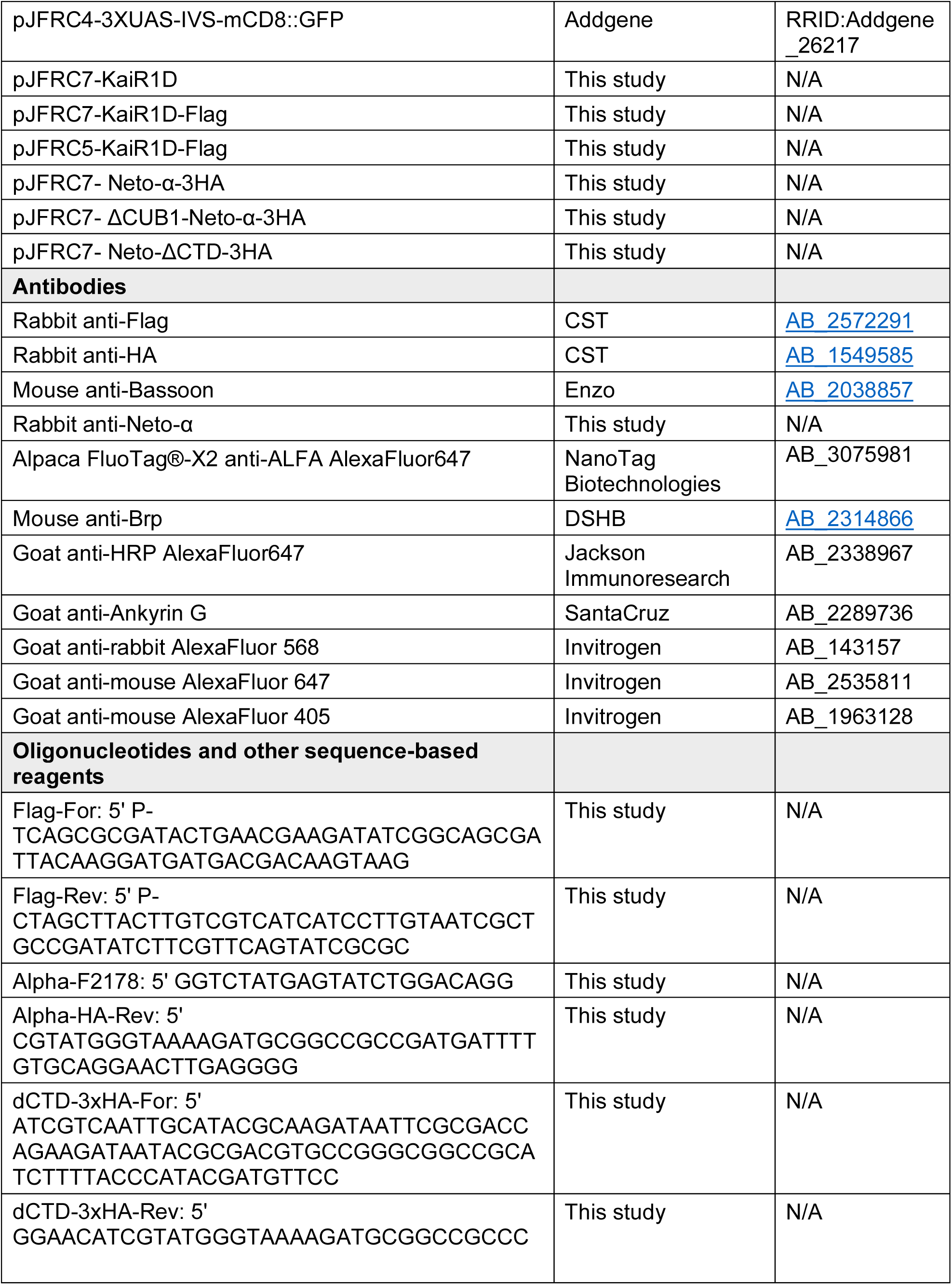
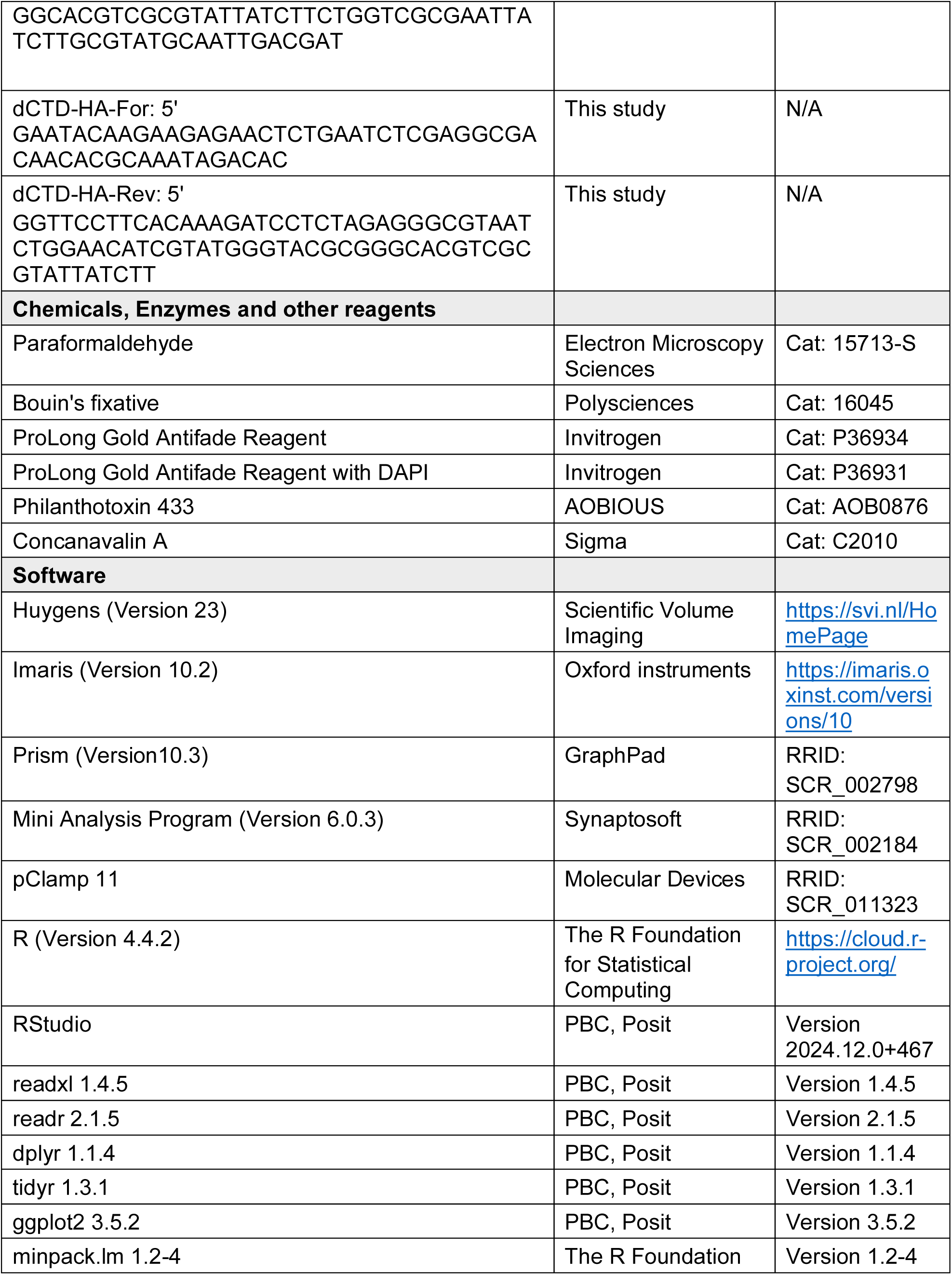
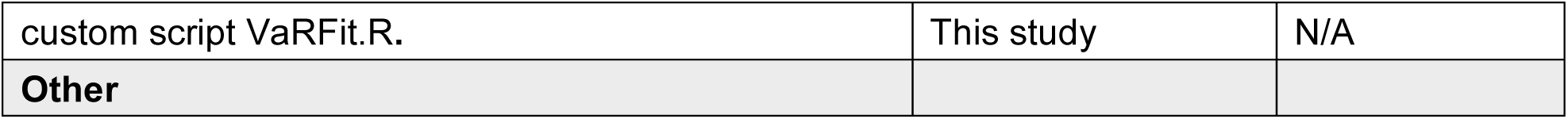

### Molecular constructs and fly stocks

The KaiR1D coding sequence was previously subcloned into a cytomegalovirus expression vector, pRK8-IRES (Li *et al*., 2016). The Flag tag was added at the C-terminal of KaiR1D (digested with BbvcI and NheI) with annealed primers:

Flag-For:

5’-P-TCAGCGCGATACTGAACGAAGATATCGGCAGCGATTACAAGGATGATGACGACAAGTAAG;

Flag-Rev:

5’ P-CTAGCTTACTTGTCGTCATCATCCTTGTAATCGCTGCCGATATCTTCGTTCAGTATCGCGC;

To generate the *UAS-KaiR1D* and *UAS-KaiR1D-Flag* transgenes, the coding sequences for KaiR1D variants were introduced downstream 20xUAS or 3xUAS in plasmids pJFRC7-20XUAS-IVS-mCD8::GFP and pJFRC4-3XUAS-IVS-mCD8::GFP respectively, gifts from Gerald Rubin (Addgene plasmids # 26220 and #26217) (Pfeiffer *et al*, 2010). These plasmids were injected at the attP docking site vk1 for phiC31 integrase-mediated transformation (Venken *et al*., 2006)

The pRK5-Neto constructs included full length Neto-α and Neto-β, derived from GH11189 and RE42119, respectively (Han *et al*., 2015), Neto-ΔCTD (M1-D478) (Ramos *et al*., 2015), and ΔCUB1 (A131/D264)-Neto-α, which was generated by QuickChange Mutagenesis (joining residues A131 and D264). To engineer the HA-tagged Neto-α variants, the end of Neto-α coding sequence was PCR amplified, joined in frame with a gene synthesized (NotI/XbaI) 3xHA cassette, and used to replace the end of Neto-α sequence in pRK5-Neto-α (Han *et al*., 2024). The ΔCUB1 variant was generated by swapping the BbvCI/BglII fragment from the untagged pRK5-ΔCUB1-Neto-α (Han *et al*., 2024). The CTD was looped out using Gibson assembly and the dCTD-3xHA set of primers. To generate the ALFA-tagged Neto-α, the 3xHA cassette was replaced with a gene synthesized NotI/XbaI fragment containing 1xALFA tag (1xAT) (Gotzke *et al*, 2019).

The series of *UAS-Neto-α-3xHA* transgenes was engineered using Gibson assembly to introduce PCR amplified (dCTD-HA-For and dCTD-HA-Rev) Neto-ΔCTD-3xHA into BglII/XbaI digested JFRC7-20XUAS-IVS-mCD8::GFP and first generate pJFRC7-Neto-ΔCTD-3xHA. The other Neto variants were next subcloned by exchanging the corresponding BglII/XbaI fragments. These plasmids were injected at the attP docking site vk14 for phiC31 integrase-mediated transformation (Venken *et al*., 2006). All molecular constructs were verified by sequencing.

The following additional fly stocks were utilized: *w^1118^*, *neto-α^null^* and *KaiR1D^null^* (Han *et al*., 2020), and *BG380-Gal4* (Budnik *et al*, 1996).

Fly stocks and genetics crosses were maintained at 25°C in an 12h light /12h dark cycle incubator. Flies were reared in vials or bottles containing Jazz-Mix food (Fisher Scientific) and analyzed at the third instar larval stage.

### *Drosophila* immunohistochemistry

Wandering third instar larvae of the desired genotypes were dissected in ice-cooled Ca^2+^-free HL-3 solution (70 mM NaCl, 5 mM KCl, 20 mM MgCl_2_, 10 mM NaHCO_3_, 5 mM trehalose, 5 mM HEPES, 115 mM sucrose) (Stewart et al., 1994; Budnik et al., 1996). The samples were fixed in 4% formaldehyde (Electron microscopy sciences) for 20 min and washed in PBS containing 0.5% Triton X-100 (PBT), incubated with primary antibodies diluted in PBT overnight at 4°C, washed with PBT, and incubated with Alexa488, Alexa568, Alexa647-conjugated secondary antibodies overnight at 4°C, washed with PBT and mounted with Prolong Gold Antifade Mountant (Invitrogen).

### Primary hippocampal neuron culture and immunohistochemistry

Hippocampal neurons were isolated from E18–19 male and female rat embryos as previously described (Sala *et al*, 2000) and seeded on 18mm-diameter coverslips coated with 3.125 μg/ml of poly–D–lysine (Corning) and 1.25 μg/ml of laminin (Thermo Fisher) at a density of ∼270 cells/mm^2^. The primary neurons were grown in Neurobasal medium (Invitrogen) supplemented with 2% B27 (Thermo Fisher) and 1% glutamax (Thermo Fisher). DIV 15 neurons were transfected with Lipofectamine 2000 (Invitrogen) and fixed at DIV 19 with 4% formaldehyde in PBS containing 4% sucrose for 15min in room temperature. Following washes with PBS, neurons were incubated with primary antibodies diluted in GBD buffer (0.18% Gelatin, 0.27% Triton X-100, 12 mM Na_2_HPO_4_, 48 mM NaH_2_PO_4_, 0.8M NaCl) overnight at 4°C, washed with PBS, and incubated with Alexa405, Alexa568, Alexa647-conjugated secondary antibodies at room temperature for 1 hour, washed with PBS and mounted with Prolong Gold (Invitrogen). The rabbit polyclonal anti-Neto-α antibodies were raised against a synthetic peptide (CKPKSGVHPLKFLHKII) (Open Biosystems) and separated by affinity purification.

### Image acquisition and analysis

For third instar larvae, each experimental replicate contained 4-5 animals per genotype that were fixed, stained and imaged together; z-stack images of muscle 4 NMJs, abdominal segments A3 and A4, were captured in the same imaging session with identical confocal settings using a laser scanning confocal microscope (CarlZeiss LSM780, 40X ApoChromat, 1.4 NA, oil immersion objective). Similarly, all hippocampal neurons within each experimental replicate were imaged with the same confocal settings. For each hippocampal neuron, we took three images: 1) a low magnification image capturing soma and dendrites and axons surrounding soma (with no zooming); 2) a high-magnification image at 4x zoom that contained primary and secondary dendrites within 100-250 μm from soma; 3) a high-magnification image at 4x zoom that contained axons >100 μm away from the soma. To image Bassoon-marked synaptic sites, we used a super-resolution confocal laser scanning microscope (CarlZeiss LSM880 Airyscan), with a 60X (1.4 NA) oil immersion objective and zoom set at 3x.

The z-stack images were deconvolved and 3D images were reconstituted using Huygens Professional software (Scientific Volume Imaging). 3D image analyses were performed using the object-to-object statistical function in Imaris (Version 10.2 Bitplane). For larval NMJ, we used the surface rendering tool to first generate a 3D model of the NMJ based on the HRP signal. We then captured the Brp and KaiR1D puncta within this NMJ volume and quantified their fluorescence intensities. For transfected hippocampal neurons, the 3D neurite volumes were similarly defined by the Venus signal. We then created surface models for Neto-α, KaiR1D, and Bassoon to identify, capture and quantify the corresponding puncta and examine their colocalization.

### *In situ* intracellular recordings

The standard larval body wall muscle preparation first developed by Jan and Jan (1976) (Jan & Jan, 1976b) was used for electrophysiological recordings (Zhang & Stewart, 2010). Wandering third instar larvae were dissected in physiological saline HL-3 saline (Stewart *et al*, 1994), washed, and immersed in HL-3 containing 0.5 mM Ca^2+^ using a custom microscope stage system (Ide, 2013). The nerve roots were cut near the exiting site of the ventral nerve cord so that the motor nerve could be picked up by a suction electrode. Intracellular recordings were made from muscle 6, abdominal segment 3 and 4. Data were used when the input resistance of the muscle was >5 MΩ and the resting membrane potential was between −60 mV and −70 mV. The input resistance of the recording microelectrode (backfilled with 3 M KCl) ranged from 20 to 25 MΩ. Muscle synaptic potentials were recorded using Axon Clamp 2B amplifier (Axon Instruments) and analyzed using pClamp 10 software. Following motor nerve stimulation with a suction electrode (200 μsec, 5 V), evoked EJPs were recorded. Six to ten EJPs evoked by low frequency of stimulation (0.1 Hz) were averaged. To calculate mEJP mean amplitudes, 50–100 events from each muscle were measured and averaged using the Mini Analysis program (Synaptosoft). Minis with a slow rise and falling time arising from neighboring electrically coupled muscle cells were excluded from analysis (Gho, 1994; Zhang *et al*, 1998). Quantal content was calculated by dividing the mean EJP by the mean mEJP after correction of EJP amplitude for nonlinear summation according to previously described methods (Feeney *et al*, 1998; Stevens, 1976). Corrected EJP amplitude = E[Ln[E/(E - recorded EJP)]], where E is the difference between reversal potential and resting potential. The reversal potential used in this correction was 0 mV (Feeney *et al*., 1998; Lagow *et al*, 2007).

### Expression of KaiR1D receptor complexes in HEK293T cells

HEK293T cells were cultured in Dulbecco’s Modified Eagle Medium (DMEM; Gibco) with 10% fetal bovine serum, 1% Glutamax and 1% Penicillin Streptomycin solution at 37 °C in a 95% oxygen and 5% carbon dioxide incubator. HEK293T cells (2 x 10^5^/ml) were plated on 5 mm diameter coverglass coated with bovine collagen (Nutragen), attached for 24h, then transfected with pRK5-based constructs of KaiR1D and Neto variants (1 µg total DNA/ ml cells) using ViaFect transfection reagent (Promega). The cells were incubated at 37°C for 3 hours, and then the temperature was decreased to 30°C. Outside-out patch-clamp recordings were performed 3 days after transfection at room temperature.

### Outside-out patch clamp recording

Outside-out patch recordings from HEK293T cells with fast solution exchange, achieved using four-bore glass tubing mounted on a P245.30 piezoelectric stack driven by a P-270 HVA amplifier (Physik Instrumente), were performed at room temperature as described previously (Horning & Mayer, 2004). Recordings were made with thin-wall borosilicate glass pipettes (resistance, 3-6 MΩ). The external solution contained (in mM) 145 NaCl, 5.4 KCl, 1.8 CaCl_2_, 1 MgCl_2_, 5 HEPES (pH 7.3). Agonist (10 mM L-glutamate) and, where indicated, 1 µM Philanthotoxin-343 (Sigma) were dissolved in external solution. The internal solution contained (in mM) 110 CsCl, 10 CsF, 0.5 CaCl_2_, 1 MgCl_2_, 10 HEPES, 5 CsBAPTA and 20 Na_2_ATP (pH 7.3). The internal solution used to test the effect of spermine contained (in mM) 140 CsCl, 10 CsF, 0.5 CaCl_2_, 1 MgCl_2_, 10 HEPES and 5 CsBAPTA (pH 7.3), with or without 0.1 spermine. The currents from outside-out patches were recorded using gap-free protocol with an Axopatch 200B amplifier (Axon Instruments), digitized with a Digidata 1550 (Molecular Devices), sampled at 20 kHz, low pass filtered at 1 kHz, and collected using pClamp10.7 (Molecular Devices). Open tip junction potentials recorded at the end of experiments typically had 10-90% rise times of < 200 µs, and data from patches for which responses to 10 mM glutamate had rise times > 400 µs were discarded. Concanavalin A (type IV), glutamate and Philanthotoxin-343 were purchased from Sigma-Aldrich.

### Outside-out patch data analysis

To calculate decay times constant of currents for various receptor channels, 50 ∼ 100 representative events from each recording were averaged and fitted using a first order exponential function for deactivation

*I(t) = I* exp*(−t/*τ)

and a double exponential function for desensitization

*I(t) = I_fast_*exp*(−t/*τ*_fast_) + I_slow_* exp*(−t/*τ*_slow_)*

where *I_x_* is the peak current amplitude of a decay component and τ_x_ is the corresponding decay time constant. To allow for easier comparison of decay times between complexes, weighted τ (ms) were calculated using the formula

*τ_w_ = (I_fast_/(I_fast_ + I_slow_))* τ_fast_ + (I_slow_/(I_fast_ + I_slow_))* τ_slow_*

For time course fitting of single-channel openings, records were filtered at 1kHz (−3dB, 8-pole Bessel), digitized at 100 kHz with a Digidata 1550 (Molecular Devices) and individual currents fitted with single channel search algorithm of pCLAMP 10.7. The histograms of open time were obtained and displayed using Clampfit software. The mean amplitude of single channel currents were determined from fits of one or two Gaussian distributions. One way ANOVA was used for statistical analysis. The data are reported as mean ± SEM.

### Non-Stationary Variance Analysis of Decay-Phase Currents

To estimate channel properties from macroscopic current traces, we implemented a custom R-based pipeline, VaRFit.R (Supplemental file) which runs in RStudio, version 2024.12.0+467, on top of R version 4.2.2. This pipeline performs non-stationary variance analysis restricted to the decay phase of responses. For each patch, individual current responses were aligned to their peak using the Mini Analysis Program (Synaptosoft, v6.03) and exported as a matrix-formatted Excel file, where each column corresponds to a single recorded event. Traces were sampled at 10 kHz. To avoid noise from the rising phase, only the post-peak decay regions of the traces were analyzed.

Specifically, the dataset was reshaped by extracting two segments, a short pre-peak interval for baseline estimation and a longer post-peak segment for variance analysis. A sliding window of 100 points was applied across the pre-stimulus region, and the interval with the lowest mean variance across all traces was selected as the baseline. Traces were excluded if their baseline variance exceeded a threshold defined as the mean plus four standard deviations across all traces. Pairwise differences between adjacent traces were computed to generate a difference matrix which was used to estimate the variance at each time point. The mean current was calculated, and variance values were corrected by subtracting the baseline variance. Both metrics were grouped into 100 equally spaced bins across the mean current range. The binned data were fitted to the parabolic equation, *Variance=a⋅I^2^+b⋅I,* using nonlinear least squares fitting (nlsLM). The number of channels was estimated as *N=-1/a*, and the unitary current as *i=b*. The open probability was calculated for each bin as *Popen=I/(i⋅N),* where *I* is the bin’s average mean current. The single-channel conductance (*γ*) was estimated using the formula *γ*=i/Vm⋅1000 (pS) assuming a holding potential of - 60mV. Plots showing the variance–mean relationship and fitted parabolas were generated using the ggplot2 package.

### Statistical analysis

All statistical analyses were performed in GraphPad Prism v10. Outliers were identified with the ROUT method (Q = 1%) and excluded prior to testing. Data are reported as mean ± standard error of the mean (SEM). Microscopy datasets were evaluated with non-parametric tests (Mann– Whitney U for two groups; Kruskal–Wallis for multiple groups with Dunn’s post hoc). Individual points denote single hippocampal neurons or larval NMJs (one or two NMJs per larvae were examined). Electrophysiology datasets were analyzed with one-way ANOVA followed by Tukey’s multiple comparisons. **** p < 0.0001; *** 0.0001 ≤ p < 0.001; ** 0.001 ≤ p < 0.01; * 0.01 ≤ p < 0.05; ns, p ≥ 0.05.

## Acknowledgments

We are grateful to Mark Mayer for continuous guidance and for comments on this manuscript. We thank members of Serpe laboratory for their feedback. W-C.H, T.H.H, R.V., P.N. and M.S. were supported by Intramural Program of the NICHD, grants ZIA HD008914 and ZIA HD008869 awarded to M.S. Z.L. was supported by the NIMH Intramural Program, grant 1ZIAMH002881. Additional funding was awarded to W-C.H. from Center from Compulsive Behavior, NIH. The contributions of the NIH authors are considered Works of the United States Government. The findings and conclusions presented in this paper are those of the authors and do not necessarily reflect the views of the NIH or the U.S. Department of Health and Human Services.

## Author contributions

W-C.H., R.V., T.H.H., P.N. and M.S. conducted the *in vivo* and *in vitro* experiments and analyzed the data. W-C.H. and Z.L. designed and conducted the *ex vivo* experiments. W-C.H., R.V., T.H.H., Z.L. and M.S. conceived the study and wrote the manuscript. All authors edited the manuscript.

## Data Availability

All data including the VaRFit.R code are available in the main figures and supplementary materials. Further information and requests for resources and reagents should be directed to the lead contact, Mihaela Serpe.

## Supplemental file

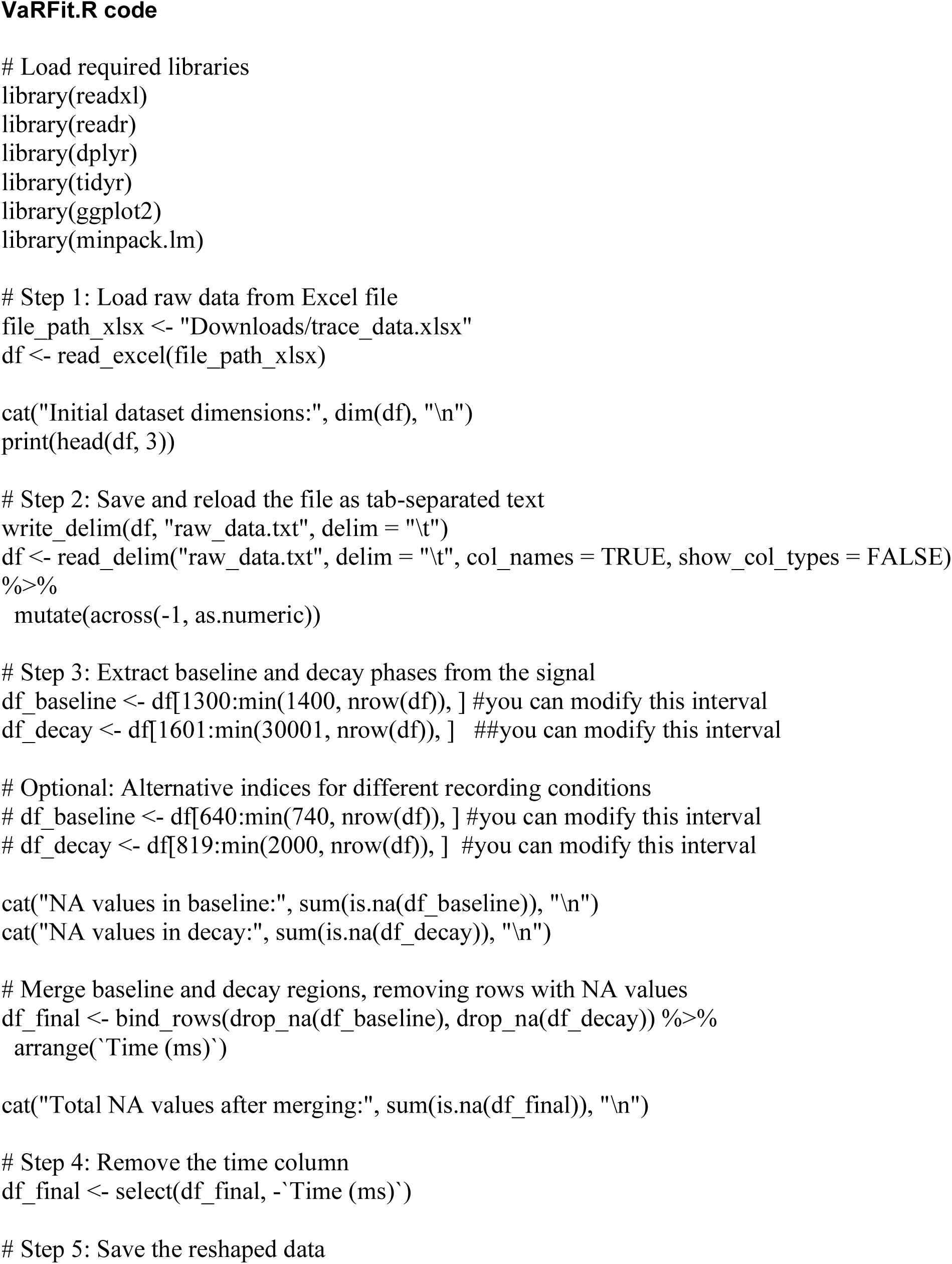
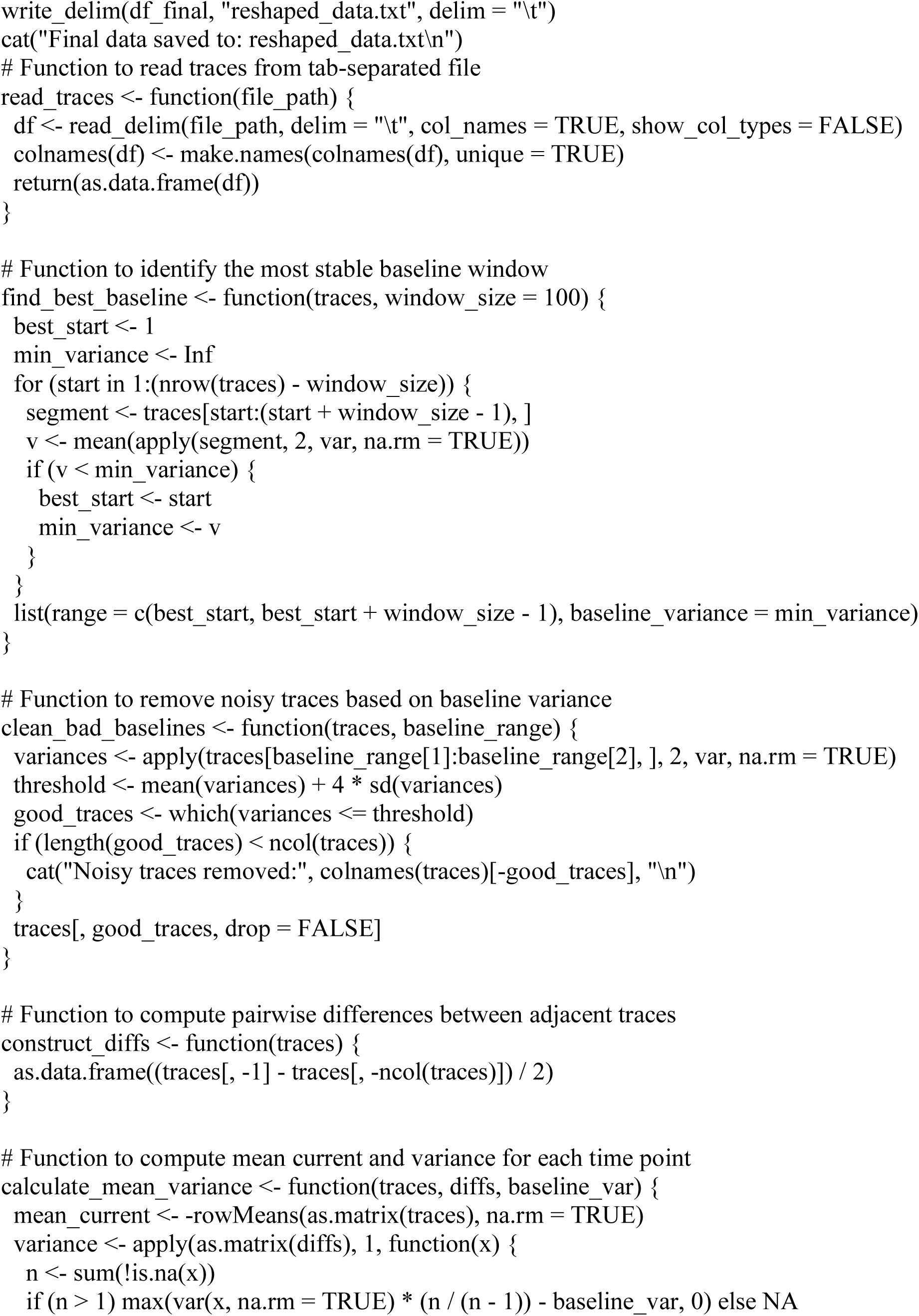
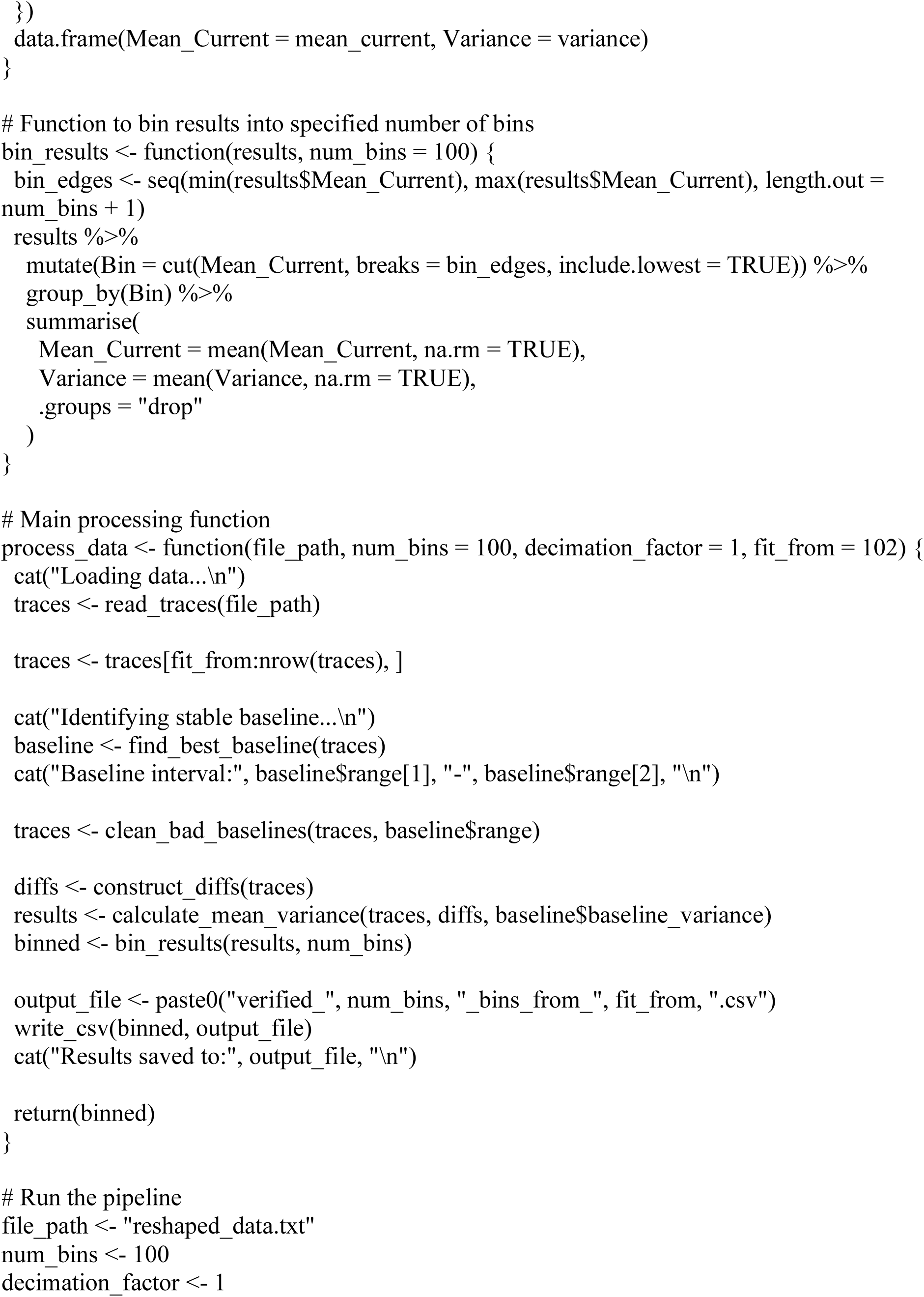
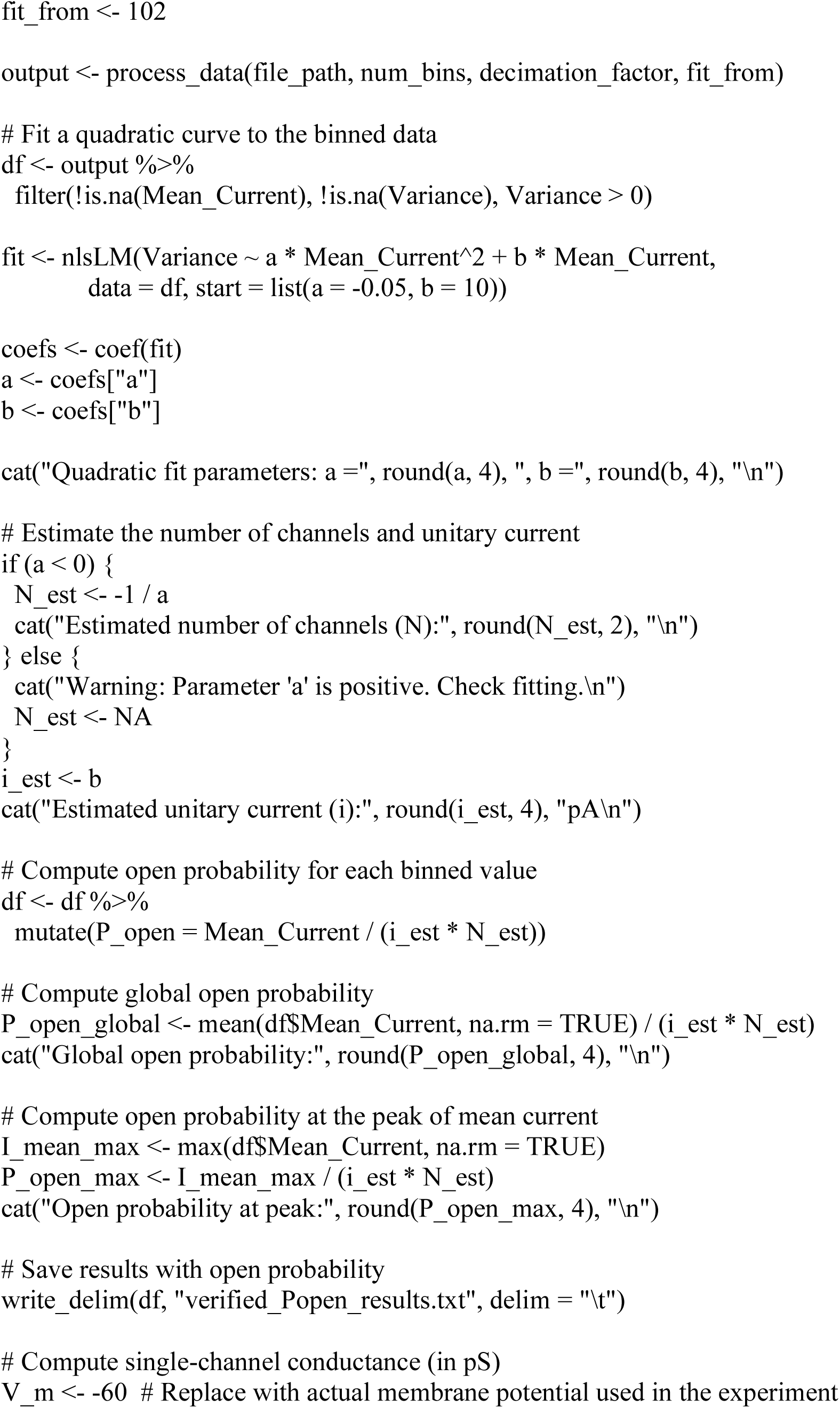
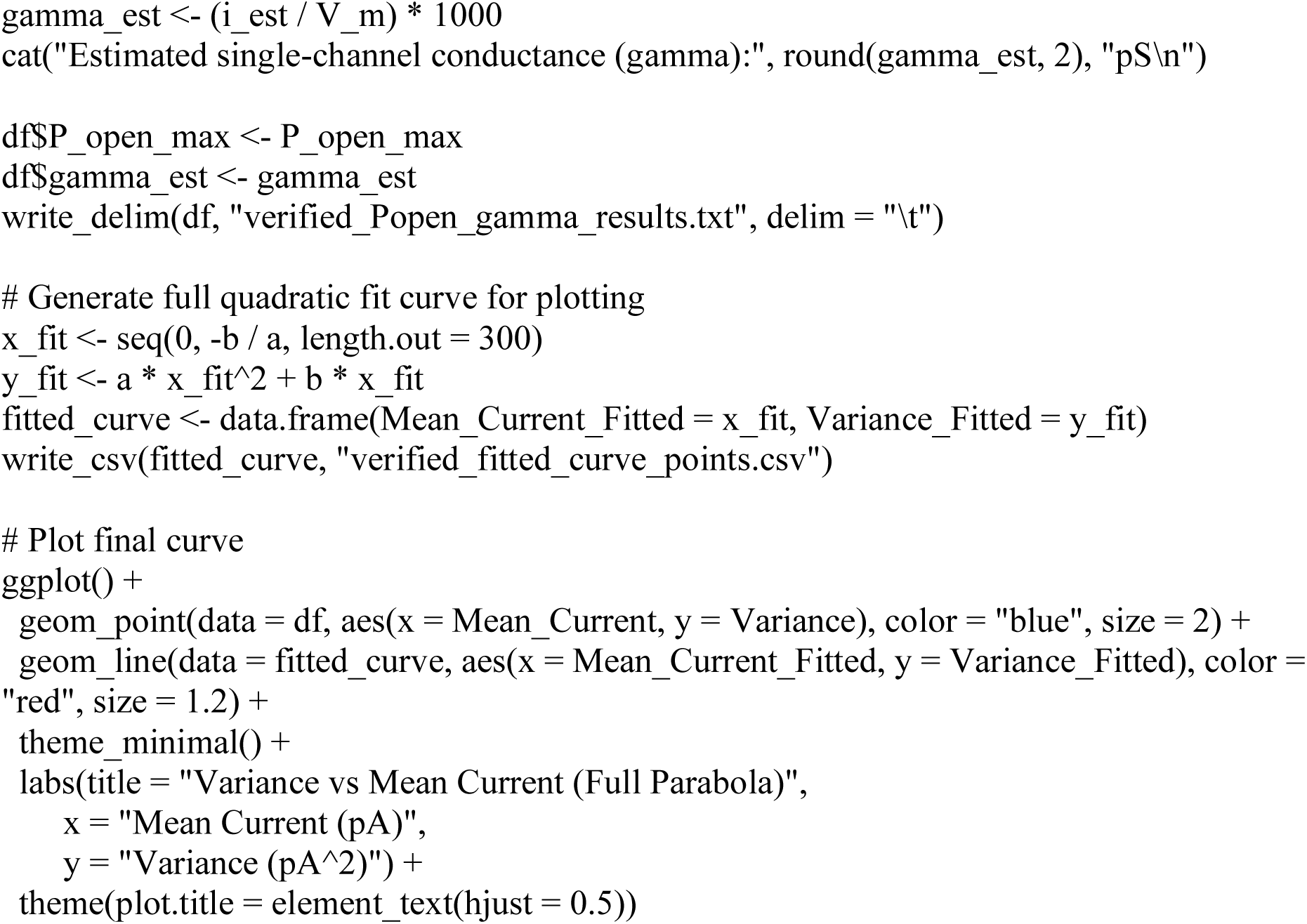

---

**Supplemental Figure S1.**
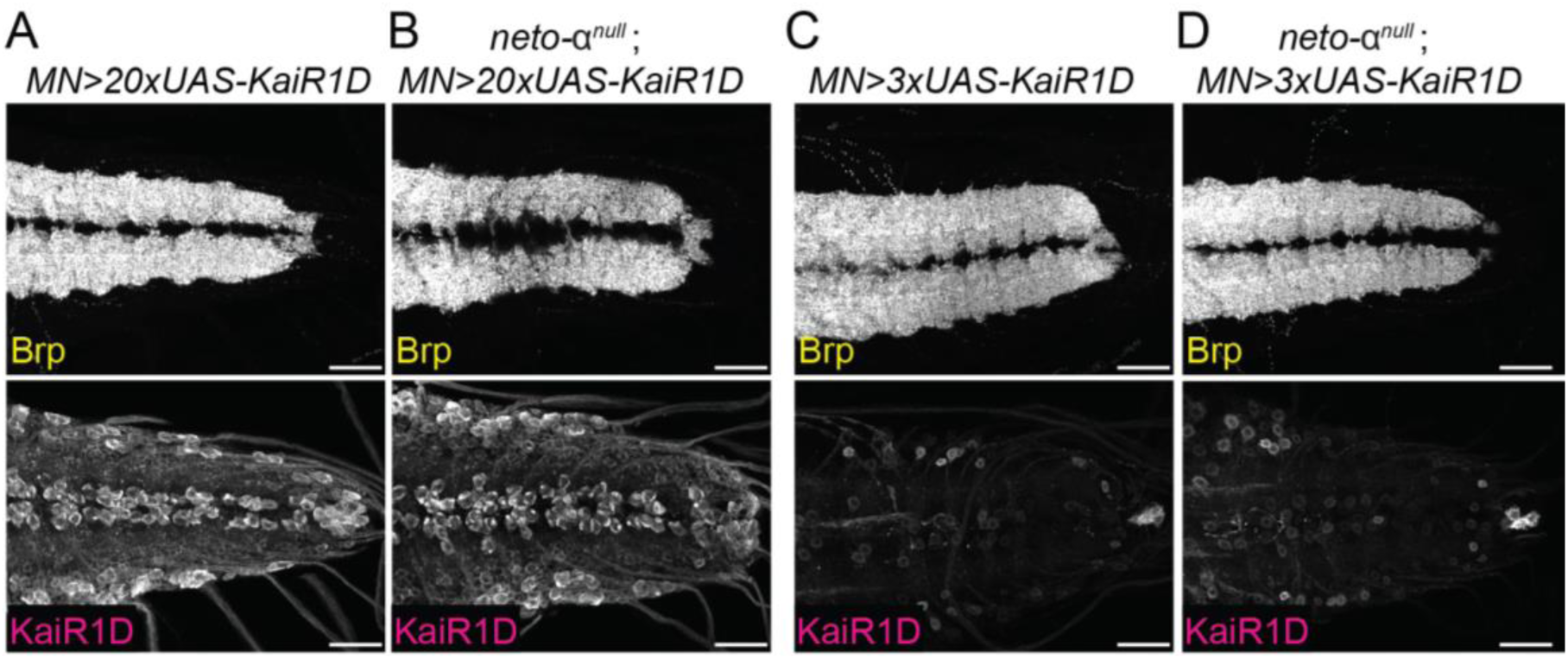
Different levels of KaiR1D expression in the ventral cord. Confocal images of third instar larva NMJ of the indicated genotypes stained for KaiR1D (magenta), Brp (yellow) and HRP (white). High levels KaiR1D expression in the motor neurons (*BG380>20xUAS-KaiR1D-Flag*) induces dramatic accumulation of KaiR1D signals in the motor neuron soma in comparison to low levels expression (*BG380>3xUAS-KaiR1D*-Flag). Note that the EJP deficits of *KaiR1D^null^* NMJs are rescued in this low expression rescue setting. Scale bars: 30 μm.

**Supplemental Figure S2.**
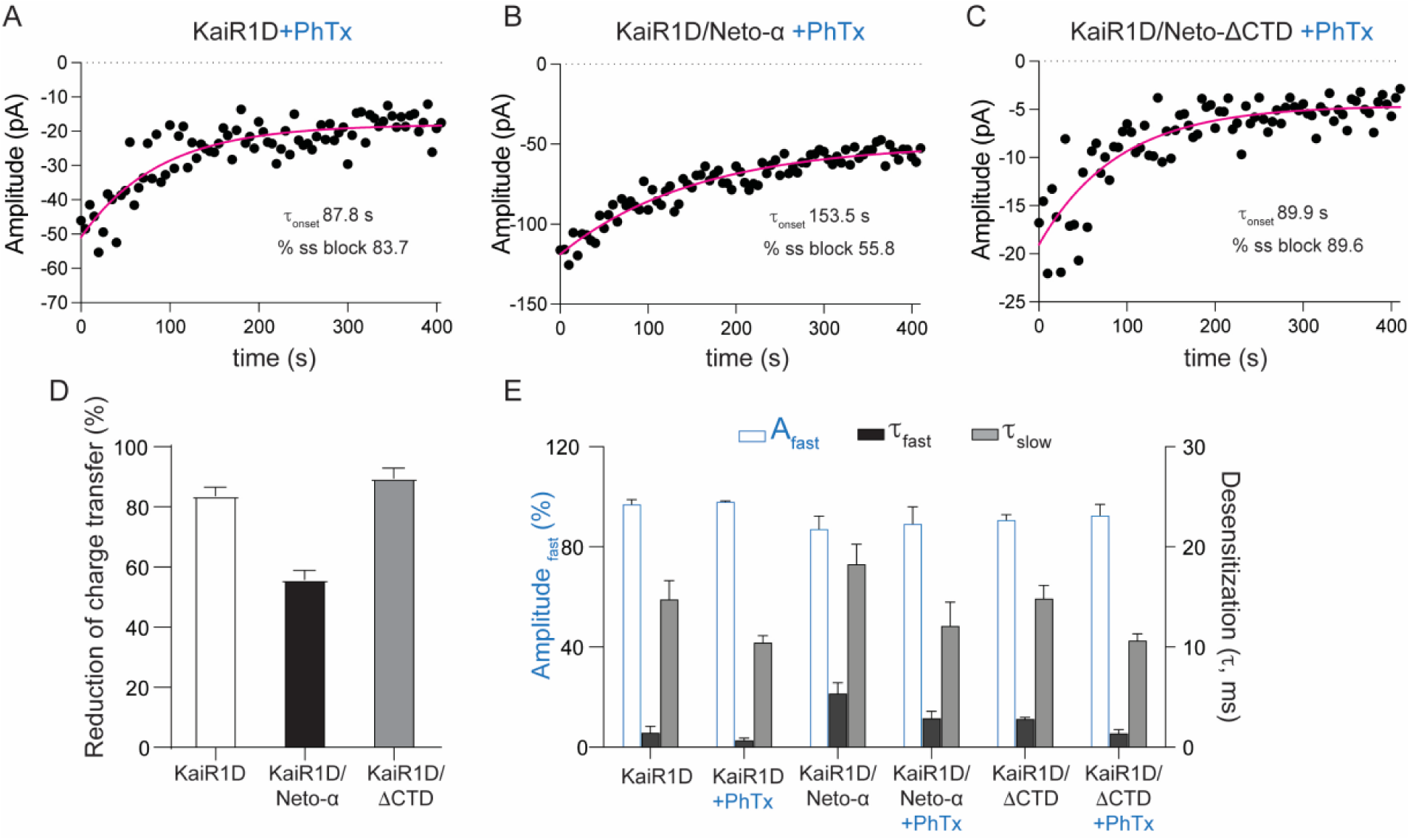
Slow onset of block by PhTx. (A-C) Data points indicate the amplitude of sequential responses to 10 mM glutamate applied for 100 ms at intervals of 1 s for KaiR1D alone (A) or together with Neto-α (B) or Neto-ΔCTD (C). The amplitude variation is due to differences in the number of channels activated from trial to trial. Red lines show single exponential fits to the decay of the response to glutamate due to onset of block by 1 µM PhTx. (D) The extent of block by PhTX at equilibrium estimated from the change in charge transfer relative to control, where a value of 100% indicates complete block. KaiR1D alone or with Neto-ΔCTD show profound block by PhTx. (E) Fits to the decay of the response to 100 ms applications of 10 mM glutamate (fit with the sum of two exponentials) before and after application of PhTx, showing mean values for %A_fast_, τ_fast_, and τ_slow_ for the indicated combinations of KaiR1D alone or together with Neto-α or Neto-ΔCTD.

**Supplemental Figure S3.**
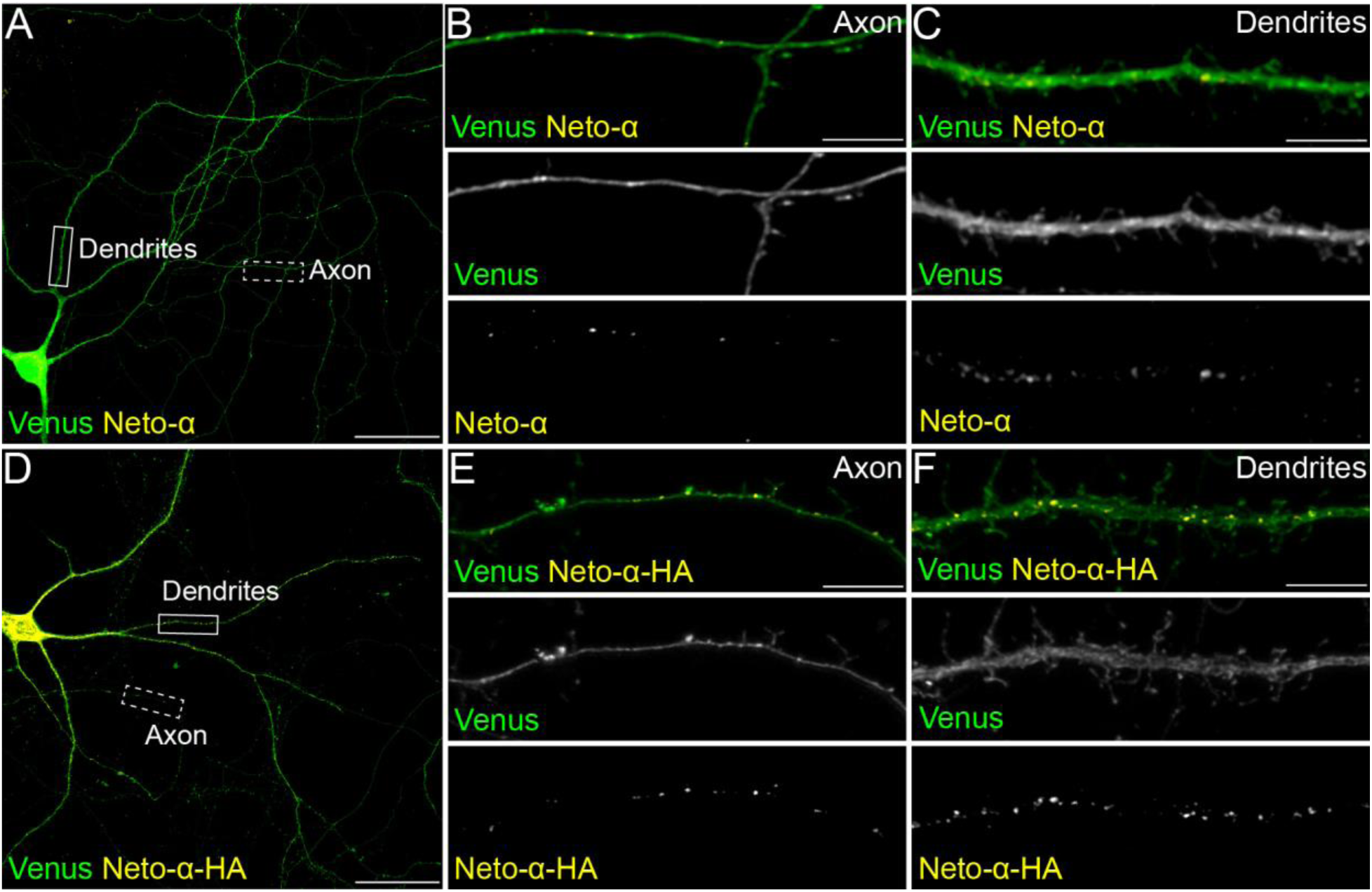
**Tagged and untagged Neto-α localize to both axons and dendrites** (A-F) Confocal images of primary rat hippocampal neurons transfected (DIV15) and stained (DIV19) for Venus (green) and Neto-α (yellow). Low magnification views (A and D) capture the cell body as well as part of the axon and dendrites, shown at higher magnification in details (B-C and E-F). Scale bars: 40 µm (A, D and G) and 5 µm (axons and dendrites details).

**Supplemental Figure S4.**
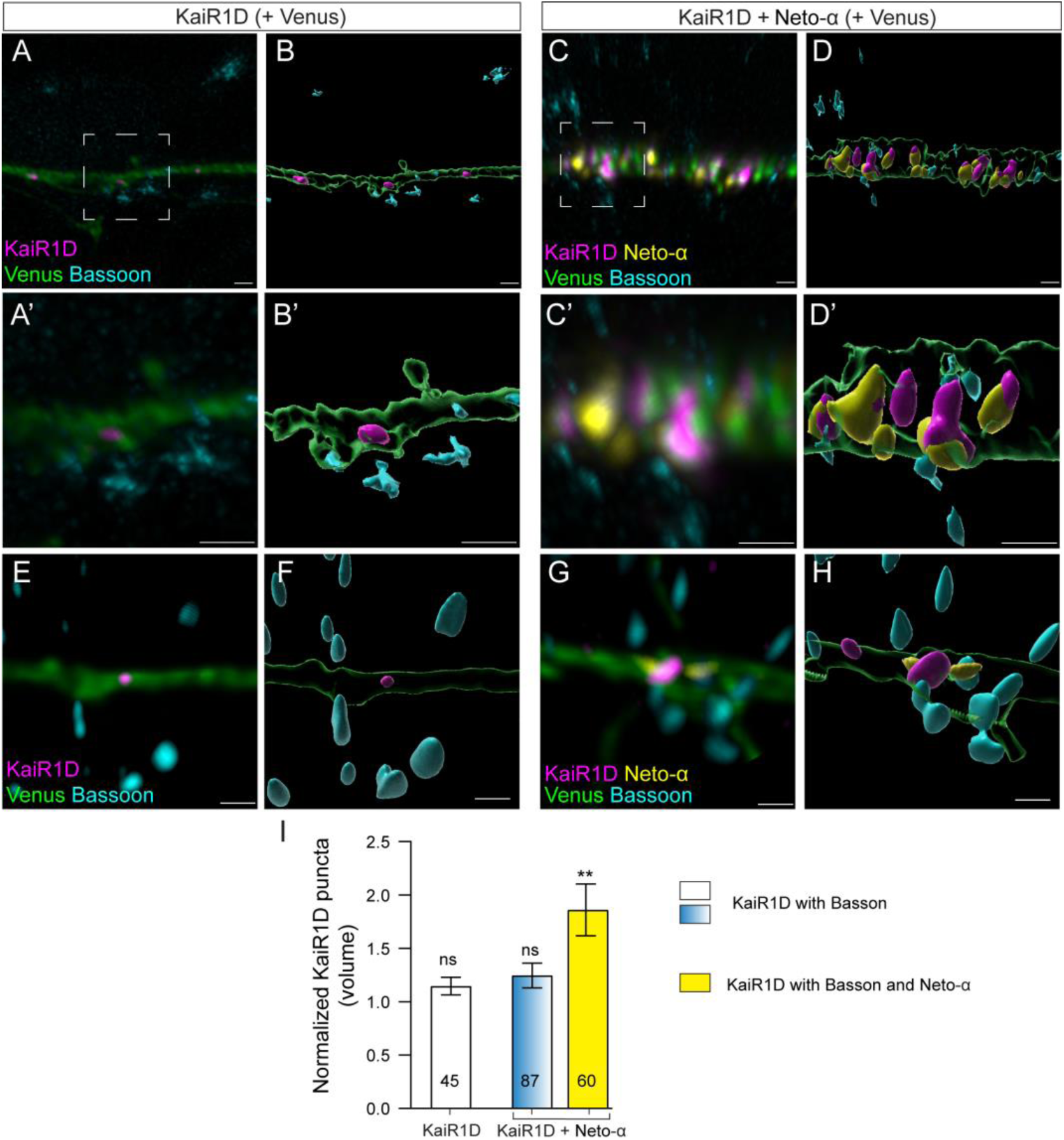
KaiR1D and Neto-α accumulate at presynaptic sites. Confocal images and 3D reconstitution of presynaptic contacts from primary rat hippocampal neurons transfected (DIV15) with KaiR1D alone (upper) or with Neto-α (lower) and stained (DIV19) for Venus (green), Bassoon (cyan), KaiR1D (magenta) and Neto-α (yellow). When transfected without Neto-α, KaiR1D rarely enters the axonal compartment and does not colocalize with Bassoon. When transfected together, KaiR1D and Neto-α they form complexes adjacent to Bassoon puncta. Data are represented as mean ± SEM. Indicated P values are from one-way ANOVA with Karuskal-Wallis multiple comparison test. **p < 0.01. Scale bar: 1 µm.

**Supplemental Figure S5.**
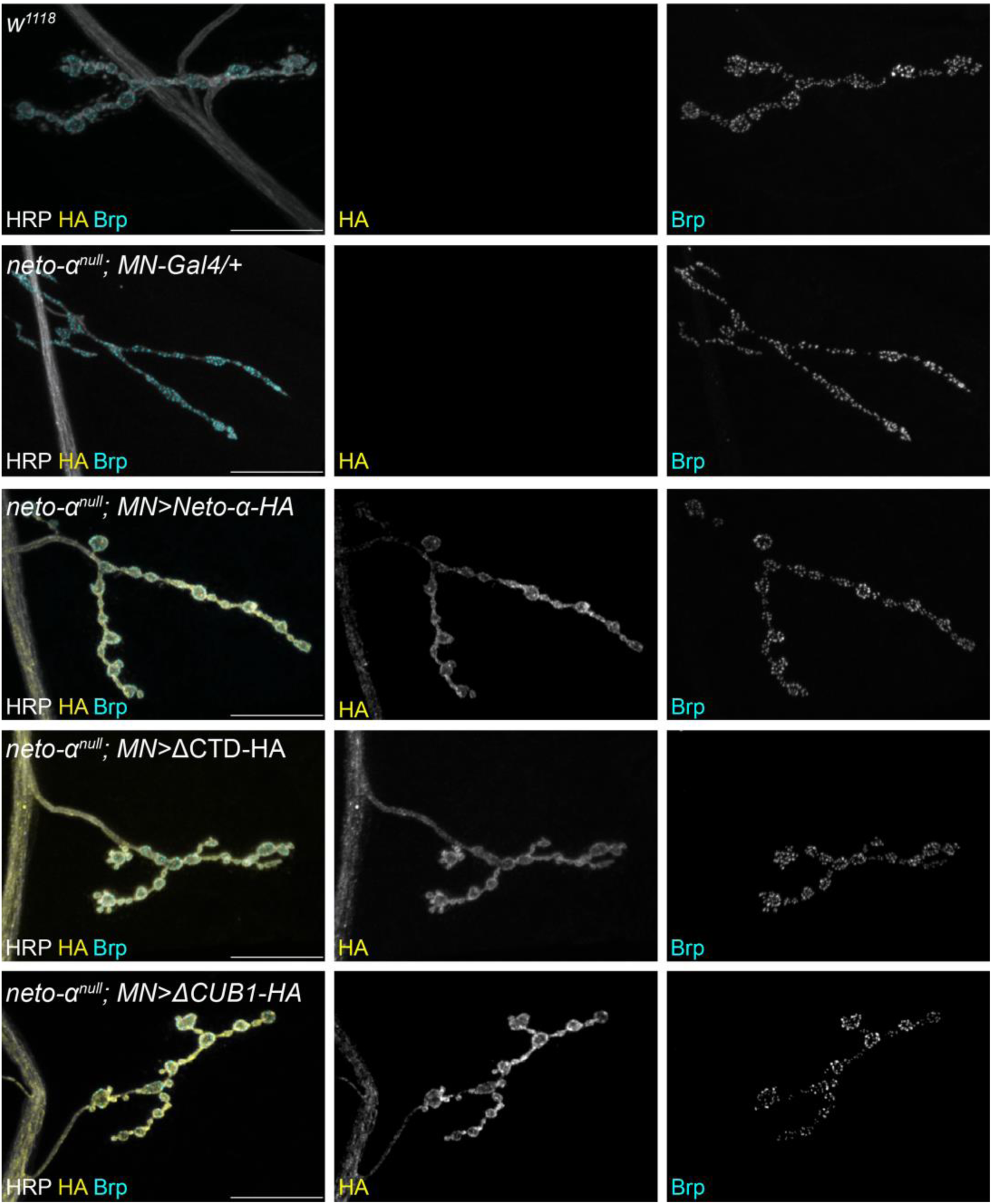
**Neto-α variants reach the presynaptic terminals in *neto-α^null^* mutants** Confocal images of third instar larva NMJ for the indicated genotypes stained for Brp (yellow), HRP (white) and various HA-tagged Neto-α variants (yellow). The first two genotypes, *w^1118^* and driver alone (*BG380-Gal4/+*) in *neto-α^null^* background, serve as controls. When expressed in the motor neurons, all three Neto-α variants examined localize to synaptic terminals, even in the absence of endogenous Neto-α. Scale bars: 20 μm.

**Supplemental Table 1:** Electrophysiological recordings related to Figure 1.

| Fig | Genotype | mEJP Amp (mV) | EJP (mV) | QC | Rin (MΩ) | V<sub>m</sub>rest (mV) | N |
| --- | --- | --- | --- | --- | --- | --- | --- |
| 1 | control | 1.15 ± 0.04 | 34.15 ± 0.97 | 30.14 ± 1.46 | 8.79 ± 0.90 | 62.36 ± 0.76 | 10 |
| 1 | *BG380-Gal4/+* | 1.03 ± 0.04 | 30.54 ± 1.01 | 29.99 ± 1.27 | 7.89 ± 0.69 | 63.53 ± 0.66 | 9 |
| 1 | *UAS-KaiR1D-Flag/+* | 1.07 ± 0.04 | 31.07 ± 0.75 | 29.43 ± 1.40 | 8.43 ± 0.42 | 64.64 ± 0.39 | 10 |
| 1 | *KaiR1D<sup>null</sup>* | 0.99 ± 0.08 | 19.94 ± 1.82 | 20.39 ± 1.82 | 13.63 ± 1.34 | 64.11 ± 1.30 | 10 |
| 1 | *KaiR1D<sup>null</sup>*<br>*BG380>KaiR1D-Flag* | 1.09 ± 0.02 | 33.21 ± 0.76 | 30.48 ± 0.57 | 6.48 ± 0.52 | 65.44 ± 0.64 | 10 |
| 1 | *BG380>KaiR1D-Flag* | 1.09 ± 0.04 | 30.28 ± 0.37 | 28.03 ± 1.16 | 8.45 ± 0.33 | 64.16 ± 0.46 | 10 |
| 1 | *neto-α<sup>null</sup>* | 1.04 ± 0.03 | 22.01 ± 0.67 | 21.29 ± 0.69 | 7.93 ± 0.46 | 66.15 ± 1.25 | 10 |
| 1 | *neto-α<sup>null</sup>*<br>*BG380>KaiR1D-Flag* | 1.06 ± 0.03 | 18.97 ± 0.24 | 17.95 ± 0.40 | 9.75 ± 0.69 | 65.36 ± 0.50 | 10 |

**Supplemental Table 2.** Kinetic analysis for currents evoked by 1 ms (deactivation) and 100 ms (desensitization) applications of 10 mM glutamate recorded using outside out patches from HEK cells transfected with the indicated KaiR1D/Neto variant combinations.

|  | Rise time<br>(ms) | Deactivation time |  | Desensitization time |  |  |  | N |
| --- | --- | --- | --- | --- | --- | --- | --- | --- |
| | | $\tau_{off}$ (ms) | A (pA) | $\tau_{fast}$ (ms) | $\tau_{slow}$ (ms) | $A_{fast}$ (%) | $\tau_w$ (ms) | |
| KaiR1D | 0.24 ± 0.01 | 0.39 ± 0.03 | 43.37 ± 12.25 | 1.07 ± 0.06 | 12.72 ± 1.61 | 89.61 ± 2.61 | 2.31 ± 0.39 | 10 |
| KaiR1D+Neto-α | 0.25 ± 0.01 | 0.39 ± 0.03 | 43.66 ± 16.37 | 4.30 ± 0.35 | 17.21 ± 1.45 | 75.89 ± 3.98 | 7.09 ± 0.34 | 10 |
| KaiR1D+Neto-β | 0.26 ± 0.01 | 0.39 ± 0.05 | 58.47 ± 18.89 | 3.90 ± 0.58 | 17.58 ± 2.21 | 91.12 ± 3.44 | 5.15 ± 0.73 | 8 |
| KaiR1D+Neto-ΔCTD | 0.25 ± 0.01 | 0.39 ± 0.03 | 17.86 ± 13.18 | 2.94 ± 0.44 | 13.25 ± 1.97 | 81.09 ± 5.80 | 4.33 ± 0.47 | 10 |
| KaiR1D+ΔCUB1-Neto-α | 0.26 ± 0.01 | 0.38 ± 0.01 | 11.77 ± 5.41 | 1.12 ± 0.22 | 21.46 ± 2.19 | 97.57 ± 0.93 | 1.68 ± 0.32 | 8 |

**Supplemental Table 3.**
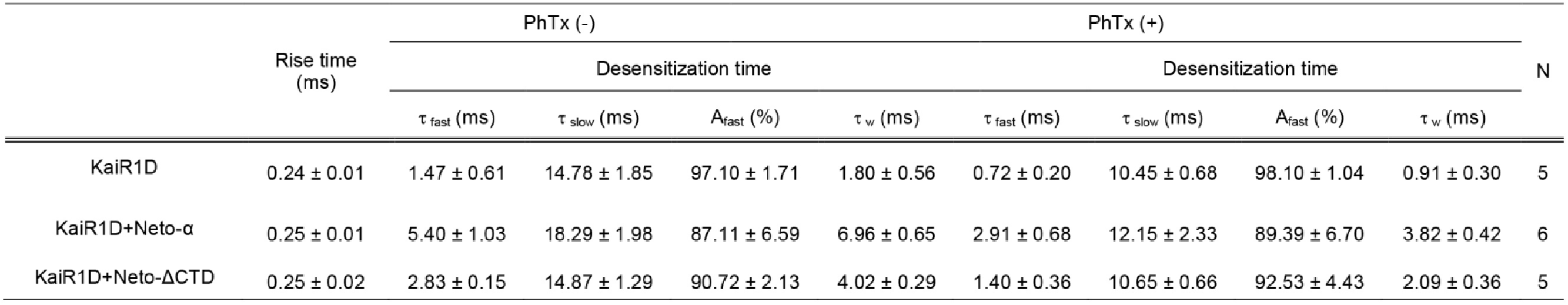
Decay kinetics for responses to 100 ms applications of 10 mM glutamate recorded from the same outside patch before and in the continuous presence of 1 µM PhTx. Responses were fit with the sum of two exponentials, revealing a 1.8 to 2-fold increase in the rate for the fast component of decay in the presence of PhTx.

**Supplemental Table 4.** Electrophysiological recordings related to Figure 7.

| Fig | Genotype | mEJP Amp (mV) | EJP (mV) | QC | Rin (MΩ) | V<sub>m</sub>rest (mV) | N |
| --- | --- | --- | --- | --- | --- | --- | --- |
| 7 | control | 1.12 ± 0.02 | 34.18 ± 0.56 | 30.48 ± 0.51 | 7.30 ± 0.36 | 65.05 ± 0.59 | 10 |
| 7 | *neto-α<sup>null</sup>* | 1.04 ± 0.03 | 22.01 ± 0.67 | 21.29 ± 0.69 | 7.93 ± 0.46 | 66.15 ± 1.25 | 10 |
| 7 | *UAS-neto-α/+* | 1.03 ± 0.05 | 30.49 ± 1.18 | 30.50 ± 2.13 | 7.89 ± 0.46 | 63.89 ± 0.55 | 10 |
| 7 | *neto-α<sup>null</sup>; MN>neto-α* | 1.11 ± 0.05 | 34.08 ± 1.37 | 31.05 ± 1.58 | 8.28 ± 0.82 | 64.13 ± 0.63 | 10 |
| 7 | *UAS-ΔCUB1-neto-α/+* | 1.02 ± 0.03 | 31.62 ± 0.36 | 31.19 ± 0.74 | 8.44 ± 0.43 | 64.45 ± 0.33 | 10 |
| 7 | *neto-α<sup>null</sup>; MN>ΔCUB1-neto-α* | 1.13 ± 0.03 | 24.37 ± 1.67 | 21.48 ± 1.41 | 9.70 ± 0.72 | 62.93 ± 0.86 | 9 |
| 7 | *MN>ΔCUB1-neto-α* | 1.05 ± 0.03 | 26.33 ± 1.18 | 25.45 ± 1.63 | 7.30 ± 0.39 | 63.97 ± 0.53 | 10 |
| 7 | *UAS-neto-ΔCTD/+* | 1.08 ± 0.04 | 31.67 ± 0.56 | 29.80 ± 1.47 | 8.35 ± 0.55 | 65.35 ± 0.62 | 9 |
| 7 | *neto-α<sup>null</sup>*<br>*MN>neto-ΔCTD* | 1.19 ± 0.24 | 33.24 ± 1.08 | 28.16 ± 1.42 | 8.76 ± 0.82 | 64.88 ± 0.41 | 9 |

## References

Bahring R, Bowie D, Benveniste M, Mayer ML (1997) Permeation and block of rat GluR6 glutamate receptor channels by internal and external polyamines. J Physiol 502 (Pt 3): 575–589

Budnik V, Koh YH, Guan B, Hartmann B, Hough C, Woods D, Gorczyca M (1996) Regulation of synapse structure and function by the Drosophila tumor suppressor gene dlg. Neuron 17: 627–640

Chaudhry C, Weston MC, Schuck P, Rosenmund C, Mayer ML (2009) Stability of ligand-binding domain dimer assembly controls kainate receptor desensitization. EMBO J 28: 1518–1530

Chittajallu R, Vignes M, Dev KK, Barnes JM, Collingridge GL, Henley JM (1996) Regulation of glutamate release by presynaptic kainate receptors in the hippocampus. Nature 379: 78–81

Contractor A, Mulle C, Swanson GT (2011) Kainate receptors coming of age: milestones of two decades of research. Trends Neurosci 34: 154–163

Copits BA, Robbins JS, Frausto S, Swanson GT (2011) Synaptic Targeting and Functional Modulation of GluK1 Kainate Receptors by the Auxiliary Neuropilin and Tolloid-Like (NETO) Proteins. J Neurosci 31: 7334–7340

Copits BA, Swanson GT (2012) Dancing partners at the synapse: auxiliary subunits that shape kainate receptor function. Nat Rev Neurosci 13: 675–686

DiAntonio A (2006) Glutamate receptors at the Drosophila neuromuscular junction. Int Rev Neurobiol 75: 165–179

Duan G-F, Ye Y, Xu S, Tao W, Zhao S, Jin T, Nicoll RA, Shi YS, Sheng N (2018) Signal peptide represses GluK1 surface and synaptic trafficking through binding to amino-terminal domain. Nature communications 9: 4879

Eldefrawi AT, Eldefrawi ME, Konno K, Mansour NA, Nakanishi K, Oltz E, Usherwood PN (1988) Structure and synthesis of a potent glutamate receptor antagonist in wasp venom. Proc Natl Acad Sci U S A 85: 4910–4913

Farias GG, Guardia CM, Britt DJ, Guo X, Bonifacino JS (2015) Sorting of Dendritic and Axonal Vesicles at the Pre-axonal Exclusion Zone. Cell reports 13: 1221–1232

Featherstone DE, Rushton E, Rohrbough J, Liebl F, Karr J, Sheng Q, Rodesch CK, Broadie K (2005) An essential Drosophila glutamate receptor subunit that functions in both central neuropil and neuromuscular junction. J Neurosci 25: 3199–3208

Feeney CJ, Karunanithi S, Pearce J, Govind CK, Atwood HL (1998) Motor nerve terminals on abdominal muscles in larval flesh flies, Sarcophaga bullata: comparisons with Drosophila. The Journal of comparative neurology 402: 197–209

Fisher JL, Mott DD (2012) The auxiliary subunits Neto1 and Neto2 reduce voltage-dependent inhibition of recombinant kainate receptors. J Neurosci 32: 12928–12933

Gho M (1994) Voltage-clamp analysis of gap junctions between embryonic muscles in Drosophila. J Physiol 481 (Pt 2): 371–383

Gotzke H, Kilisch M, Martinez-Carranza M, Sograte-Idrissi S, Rajavel A, Schlichthaerle T, Engels N, Jungmann R, Stenmark P, Opazo F, Frey S (2019) The ALFA-tag is a highly versatile tool for nanobody-based bioscience applications. Nature communications 10: 4403

Griffith TN, Swanson GT (2015) Identification of critical functional determinants of kainate receptor modulation by auxiliary protein Neto2. J Physiol 593: 4815–4833

Han TH, Dharkar P, Mayer ML, Serpe M (2015) Functional reconstitution of Drosophila melanogaster NMJ glutamate receptors. Proc Natl Acad Sci U S A 112: 6182–6187

Han TH, Vicidomini R, Ramos CI, Mayer ML, Serpe M (2024) The gating properties of Drosophila NMJ glutamate receptors and their dependence on Neto. J Physiol 602: 7043–7064

Han TH, Vicidomini R, Ramos CI, Wang Q, Nguyen P, Jarnik M, Lee CH, Stawarski M, Hernandez RX, Macleod GT, Serpe M (2020) Neto-alpha Controls Synapse Organization and Homeostasis at the Drosophila Neuromuscular Junction. Cell reports 32: 107866

Hansen KB, Wollmuth LP, Bowie D, Furukawa H, Menniti FS, Sobolevsky AI, Swanson GT, Swanger SA, Greger IH, Nakagawa T et al (2021) Structure, Function, and Pharmacology of Glutamate Receptor Ion Channels. Pharmacol Rev 73: 298–487

He L, Sun J, Gao Y, Li B, Wang Y, Dong Y, An W, Li H, Yang B, Ge Y et al (2021) Kainate receptor modulation by NETO2. Nature 599: 325–329

Higuchi M, Single FN, Kohler M, Sommer B, Sprengel R, Seeburg PH (1993) RNA editing of AMPA receptor subunit GluR-B: a base-paired intron-exon structure determines position and efficiency. Cell 75: 1361–1370

Horning MS, Mayer ML (2004) Regulation of AMPA receptor gating by ligand binding core dimers. Neuron 41: 379–388

Huettner JE (1990) Glutamate receptor channels in rat DRG neurons: activation by kainate and quisqualate and blockade of desensitization by Con A. Neuron 5: 255–266

Ide D (2013) Electrophysiology tool construction. Current protocols in neuroscience / editorial board, Jacqueline N Crawley [et al] Chapter 6: Unit 6 26

Jackson AC, Nicoll RA (2011) The expanding social network of ionotropic glutamate receptors: TARPs and other transmembrane auxiliary subunits. Neuron 70: 178–199

Jan LY, Jan YN (1976a) L-glutamate as an excitatory transmitter at the Drosophila larval neuromuscular junction. J Physiol 262: 215–236

Jan LY, Jan YN (1976b) Properties of the larval neuromuscular junction in Drosophila melanogaster. J Physiol 262: 189–214

Karst H, Piek T (1991) Structure-activity relationship of philanthotoxins--II. Effects on the glutamate gated ion channels of the locust muscle fibre membrane. Comp Biochem Physiol C 98: 479–489

Karuppudurai T, Lin TY, Ting CY, Pursley R, Melnattur KV, Diao F, White BH, Macpherson LJ, Gallio M, Pohida T, Lee CH (2014) A hard-wired glutamatergic circuit pools and relays UV signals to mediate spectral preference in Drosophila. Neuron 81: 603–615

Kim YJ, Bao H, Bonanno L, Zhang B, Serpe M (2012) Drosophila Neto is essential for clustering glutamate receptors at the neuromuscular junction. Genes Dev 26: 974–987

Kim YJ, Igiesuorobo O, Ramos CI, Bao H, Zhang B, Serpe M (2015) Prodomain removal enables neto to stabilize glutamate receptors at the Drosophila neuromuscular junction. PLoS genetics 11: e1004988

Kiragasi B, Wondolowski J, Li Y, Dickman DK (2017) A Presynaptic Glutamate Receptor Subunit Confers Robustness to Neurotransmission and Homeostatic Potentiation. Cell reports 19: 2694–2706

Kittel RJ, Wichmann C, Rasse TM, Fouquet W, Schmidt M, Schmid A, Wagh DA, Pawlu C, Kellner RR, Willig KI et al (2006) Bruchpilot promotes active zone assembly, Ca2+ channel clustering, and vesicle release. Science 312: 1051–1054

Kohler M, Burnashev N, Sakmann B, Seeburg PH (1993) Determinants of Ca2+ permeability in both TM1 and TM2 of high affinity kainate receptor channels: diversity by RNA editing. Neuron 10: 491–500

Lagow RD, Bao H, Cohen EN, Daniels RW, Zuzek A, Williams WH, Macleod GT, Sutton RB, Zhang B (2007) Modification of a hydrophobic layer by a point mutation in syntaxin 1A regulates the rate of synaptic vesicle fusion. PLoS Biol 5: e72

Lerma J, Marques JM (2013) Kainate receptors in health and disease. Neuron 80: 292–311

Li Y, Dharkar P, Han TH, Serpe M, Lee CH, Mayer ML (2016) Novel Functional Properties of Drosophila CNS Glutamate Receptors. Neuron 92: 1036–1048

Li YJ, Duan GF, Sun JH, Wu D, Ye C, Zang YY, Chen GQ, Shi YY, Wang J, Zhang W, Shi YS (2019) Neto proteins regulate gating of the kainate-type glutamate receptor GluK2 through two binding sites. J Biol Chem 294: 17889–17902

Lomash RM, Sheng N, Li Y, Nicoll RA, Roche KW (2017) Phosphorylation of the kainate receptor (KAR) auxiliary subunit Neto2 at serine 409 regulates synaptic targeting of the KAR subunit GluK1. J Biol Chem 292: 15369–15377

Lomash S, Chittori S, Brown P, Mayer ML (2013) Anions mediate ligand binding in Adineta vaga glutamate receptor ion channels. Structure 21: 414–425

MacDermott AB, Role LW, Siegelbaum SA (1999) Presynaptic ionotropic receptors and the control of transmitter release. Annual review of neuroscience 22: 443–485

Marrus SB, Portman SL, Allen MJ, Moffat KG, DiAntonio A (2004) Differential localization of glutamate receptor subunits at the Drosophila neuromuscular junction. J Neurosci 24: 1406–1415

Matsuda K (2017) Synapse organization and modulation via C1q family proteins and their receptors in the central nervous system. Neuroscience research 116: 46–53

Moroz LL, Nikitin MA, Policar PG, Kohn AB, Romanova DY (2021) Evolution of glutamatergic signaling and synapses. Neuropharmacology 199: 108740

Orav E, Atanasova T, Shintyapina A, Kesaf S, Kokko M, Partanen J, Taira T, Lauri SE (2017) NETO1 Guides Development of Glutamatergic Connectivity in the Hippocampus by Regulating Axonal Kainate Receptors. eNeuro 4

Orvis J, Albertin CB, Shrestha P, Chen S, Zheng M, Rodriguez CJ, Tallon LJ, Mahurkar A, Zimin AV, Kim M et al (2022) The evolution of synaptic and cognitive capacity: Insights from the nervous system transcriptome of Aplysia. Proc Natl Acad Sci U S A 119: e2122301119

Palacios-Filardo J, Aller MI, Lerma J (2016) Synaptic Targeting of Kainate Receptors. Cereb Cortex 26: 1464–1472

Panchenko VA, Glasser CR, Partin KM, Mayer ML (1999) Amino acid substitutions in the pore of rat glutamate receptors at sites influencing block by polyamines. J Physiol 520 Pt 2: 337–357

Pfeiffer BD, Ngo TT, Hibbard KL, Murphy C, Jenett A, Truman JW, Rubin GM (2010) Refinement of tools for targeted gene expression in Drosophila. Genetics 186: 735–755

Pinheiro PS, Mulle C (2008) Presynaptic glutamate receptors: physiological functions and mechanisms of action. Nat Rev Neurosci 9: 423–436

Pinheiro PS, Perrais D, Coussen F, Barhanin J, Bettler B, Mann JR, Malva JO, Heinemann SF, Mulle C (2007) GluR7 is an essential subunit of presynaptic kainate autoreceptors at hippocampal mossy fiber synapses. Proc Natl Acad Sci U S A 104: 12181–12186

Piyankarage SC, Augustin H, Grosjean Y, Featherstone DE, Shippy SA (2008) Hemolymph amino acid analysis of individual Drosophila larvae. Anal Chem 80: 1201–1207

Poulsen MH, Poshtiban A, Klippenstein V, Ghisi V, Plested AJR (2019) Gating modules of the AMPA receptor pore domain revealed by unnatural amino acid mutagenesis. Proc Natl Acad Sci U S A 116: 13358–13367

Qin G, Schwarz T, Kittel RJ, Schmid A, Rasse TM, Kappei D, Ponimaskin E, Heckmann M, Sigrist SJ (2005) Four different subunits are essential for expressing the synaptic glutamate receptor at neuromuscular junctions of Drosophila. J Neurosci 25: 3209–3218

Ramos CI, Igiesuorobo O, Wang Q, Serpe M (2015) Neto-mediated intracellular interactions shape postsynaptic composition at the Drosophila neuromuscular junction. PLoS genetics 11: e1005191

Ramos-Vicente D, Bayes A (2020) AMPA receptor auxiliary subunits emerged during early vertebrate evolution by neo/subfunctionalization of unrelated proteins. Open Biol 10: 200234

Ramos-Vicente D, Grant SG, Bayes A (2021) Metazoan evolution and diversity of glutamate receptors and their auxiliary subunits. Neuropharmacology 195: 108640

Sala C, Rudolph-Correia S, Sheng M (2000) Developmentally regulated NMDA receptor-dependent dephosphorylation of cAMP response element-binding protein (CREB) in hippocampal neurons. J Neurosci 20: 3529–3536

Schmitz D, Mellor J, Frerking M, Nicoll RA (2001) Presynaptic kainate receptors at hippocampal mossy fiber synapses. Proc Natl Acad Sci U S A 98: 11003–11008

Stevens CF (1976) A comment on Martin’s relation. Biophysical journal 16: 891–895

Stewart BA, Atwood HL, Renger JJ, Wang J, Wu CF (1994) Improved stability of Drosophila larval neuromuscular preparations in haemolymph-like physiological solutions. J Comp Physiol A 175: 179–191

Straub C, Tomita S (2012) The regulation of glutamate receptor trafficking and function by TARPs and other transmembrane auxiliary subunits. Curr Opin Neurobiol 22: 488–495

Stroebel D, Paoletti P (2021) Architecture and function of NMDA receptors: an evolutionary perspective. J Physiol 599: 2615–2638

Takeuchi A, Takeuchi N (1964) The Effect on Crayfish Muscle of Iontophoretically Applied Glutamate. The Journal of physiology 170: 296–317

Tang M, Pelkey KA, Ng D, Ivakine E, McBain CJ, Salter MW, McInnes RR (2011) Neto1 Is an Auxiliary Subunit of Native Synaptic Kainate Receptors. J Neurosci 31: 10009–10018

tom Dieck S, Sanmarti-Vila L, Langnaese K, Richter K, Kindler S, Soyke A, Wex H, Smalla KH, Kampf U, Franzer JT et al (1998) Bassoon, a novel zinc-finger CAG/glutamine-repeat protein selectively localized at the active zone of presynaptic nerve terminals. J Cell Biol 142: 499–509

Tomita S, Castillo PE (2012) Neto1 and Neto2: auxiliary subunits that determine key properties of native kainate receptors. J Physiol 590: 2217–2223

Venken KJ, He Y, Hoskins RA, Bellen HJ (2006) P[acman]: a BAC transgenic platform for targeted insertion of large DNA fragments in D. melanogaster. Science 314: 1747–1751

Wagh DA, Rasse TM, Asan E, Hofbauer A, Schwenkert I, Durrbeck H, Buchner S, Dabauvalle MC, Schmidt M, Qin G et al (2006) Bruchpilot, a protein with homology to ELKS/CAST, is required for structural integrity and function of synaptic active zones in Drosophila. Neuron 49: 833–844

Wong LA, Mayer ML (1993) Differential modulation by cyclothiazide and concanavalin A of desensitization at native alpha-amino-3-hydroxy-5-methyl-4-isoxazolepropionic acid- and kainate-preferring glutamate receptors. Mol Pharmacol 44: 504–510

Wyeth MS, Pelkey KA, Yuan X, Vargish G, Johnston AD, Hunt S, Fang C, Abebe D, Mahadevan V, Fisahn A et al (2017) Neto Auxiliary Subunits Regulate Interneuron Somatodendritic and Presynaptic Kainate Receptors to Control Network Inhibition. Cell reports 20: 2156–2168

Xu J, Marshall JJ, Fernandes HB, Nomura T, Copits BA, Procissi D, Mori S, Wang L, Zhu Y, Swanson GT, Contractor A (2017) Complete Disruption of the Kainate Receptor Gene Family Results in Corticostriatal Dysfunction in Mice. Cell reports 18: 1848–1857

Zhang B, Koh YH, Beckstead RB, Budnik V, Ganetzky B, Bellen HJ (1998) Synaptic vesicle size and number are regulated by a clathrin adaptor protein required for endocytosis. Neuron 21: 1465–1475

Zhang B, Stewart B (2010) Electrophysiological recording from Drosophila larval body-wall muscles. Cold Spring Harb Protoc 2010: pdb prot5487

Zhang W, St-Gelais F, Grabner CP, Trinidad JC, Sumioka A, Morimoto-Tomita M, Kim KS, Straub C, Burlingame AL, Howe JR, Tomita S (2009) A transmembrane accessory subunit that modulates kainate-type glutamate receptors. Neuron 61: 385–396

Zhang Y, Nayeem N, Nanao MH, Green T (2006) Interface interactions modulating desensitization of the kainate-selective ionotropic glutamate receptor subunit GluR6. J Neurosci 26: 10033–10042

